# Nuclear Hormone Receptor NHR-49/HNF4α Couples Fertility Regulation to Resource Allocation and Longevity in *C. elegans*

**DOI:** 10.1101/2025.07.14.664468

**Authors:** Sharada Gopal, Amaresh Chaturbedi, Tej Ramachandrula, John OuYang, Rebecca Rodell, Siu Sylvia Lee

## Abstract

The nuclear hormone receptor NHR-49, a homolog of mammalian PPARα and HNF4α, is a key transcriptional regulator of nutrition sensing and fatty acid metabolism in *Caenorhabditis elegans*. Here we uncovered a new function of NHR-49 in reproduction - controlling oocyte activation and ovulation. Loss of NHR-49 causes inappropriate oocyte activation and laying of unfertilized oocytes in the absence of sperm, resulting in rapid loss of yolk and stored fat, and drastically shortening of lifespan. We further demonstrated that prevention of yolk transfer into the oocytes largely restore fat storage and partially rescue lifespan in the *nhr-49* mutants. Additionally, NHR-49 appears to couple germline proliferation to nutritional status, as evidenced by its requirement for starvation-induced reduction in germline proliferation. Mechanistically, we showed that NHR-49 primarily acts in somatic cells, rather than the germline itself, to regulate oocyte activation and ovulation. We further demonstrated that NHR-49 binds to the promoter of GSA-1 and may stimulate its expression. GSA-1 encodes a G-protein coupled receptor known to act in the gonadal sheath cells to couple sperm sensing and oocyte activation. Our findings therefore suggest a model whereby NHR-49 regulates the expression of GSA-1, which in turn regulates oocyte activation in response to sperm signal. Overall, our findings suggest a mechanistic link between nutrition sensing and fertility and point to regulated retention of reproductive resources to be critical for maintaining longevity.

**Highlights:**

- A new role of NHR-49 in regulating oocyte activation and ovulation
- Inappropriate laying of unfertilized oocytes contributes to the fat loss and lifespan shortening of *nhr-49* mutants
- NHR-49 acts from somatic cells, not germline, to restrain oocyte activation
- NHR-49 binds to the promoter of GSA-1 and likely represses *gsa-1* expression to regulate oocyte activation

Graphical Abstract:
NHR-49 represses GSA-1 in sheath cells to regulate oocyte activation: We propose that NHR-49 responds to nutritional cues and represses the expression of GSA-1 in sheath cells, which results in aberrant oocyte activation and ovulation. This leads to yolk/fat loss and shortening of parental lifespan.

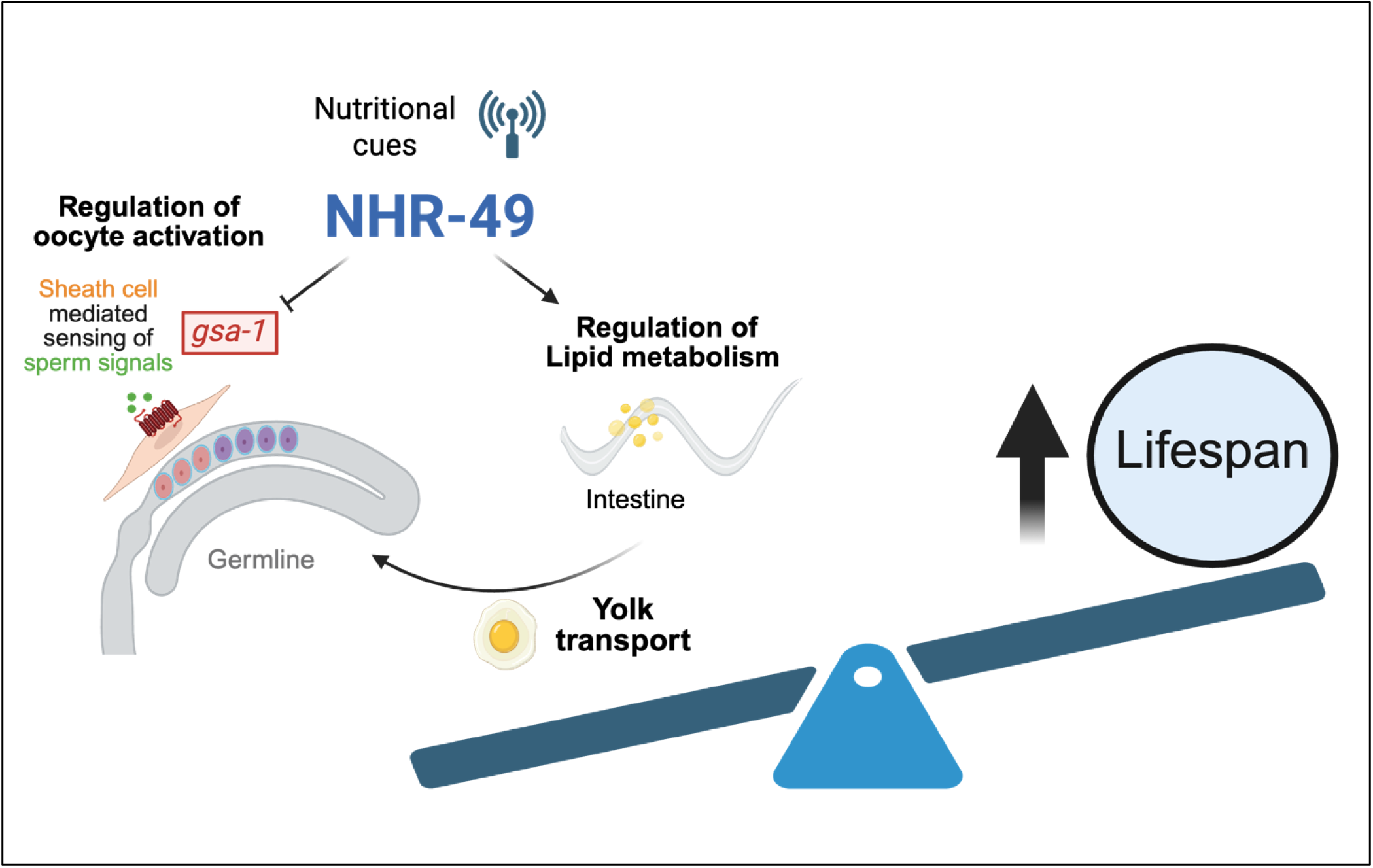

## Introduction

Reproduction is one of the most energetically demanding biological processes^1^, requiring organisms to tightly coordinate reproductive activity with metabolic status^2,3^. In nutrient-poor conditions, animals often suppress germline proliferation, delay ovulation, or halt reproduction entirely to conserve resources^4^. These adjustments are especially crucial for optimizing reproductive success and ensuring the survival of both parent and offspring^5–8^. However, the molecular mechanisms that link nutritional cues to reproductive timing remain incompletely understood.

*C. elegans* are primarily hermaphrodites and possess both oocytes and sperm, making them capable of self-fertilization^9^. During the larval stage 4 (L4), the germline stem cells in the distal region actively proliferate, and the cells further away from the distal tip cell undergo meiosis, where they take up the sperm fate and the developed sperm are stored in the spermatheca^10^. From the young adult stage onwards, germ cells switch fate to an oogenic fate and the developing oocytes are arrested in meiosis I and stored in the oviduct^11^. A highly regulated process enables the oocytes most proximal to the spermatheca to sense the presence of sperm, become activated, i.e. begin to progress pass meiosis I, and simultaneously be ovulated into the spermatheca where they become fertilized; the fertilized embryos continue to develop in the uterus until they are laid outside of the parent at around 300 cells stage^12^. The FEM (feminization) and FOG (feminization of germline) genes are crucial regulators of sex determination and spermatogenesis in *C. elegans*^1^. Loss-of-function mutations of *fem-3*, *fog-2* or *fog-3* result in defective spermatogenesis and feminization of worms that are incapable of self-fertilization ^13^**(Fig. 1A)**. Similar to hermaphrodites, the germline in these mutants contain mitotic and meiotic cells; However, the germ cells do not take up the spermatogenic fate during development. This results in an organism that has no sperm in their spermatheca. In the absence of males, these mutants accumulate ∼20 oocytes in their oviduct and the process of germline stem cell proliferation and differentiation appear paused. Upon mating with males, or when sperm are provided ^14^, the feminized mutants undergo normal stem cell proliferation and differentiation into oocytes, which are fertilized by the sperm. Mated feminized mutants produce similar number of healthy progeny as for mated hermaphrodites. Thus, feminized mutants in *C. elegans* resemble females of non-hermaphroditic species^14^. Feminized *C. elegans* mutants have been crucial in revealing the molecular pathways that regulate how oocytes sense sperm and become activated and ovulated^15–17^. Interestingly, we recently reported that unmated feminized mutants store elevated levels of fat and exhibit extended lifespan, likely due to a lack of progeny production and consequent reduced energy expenditure^18^.

**Figure 1.**
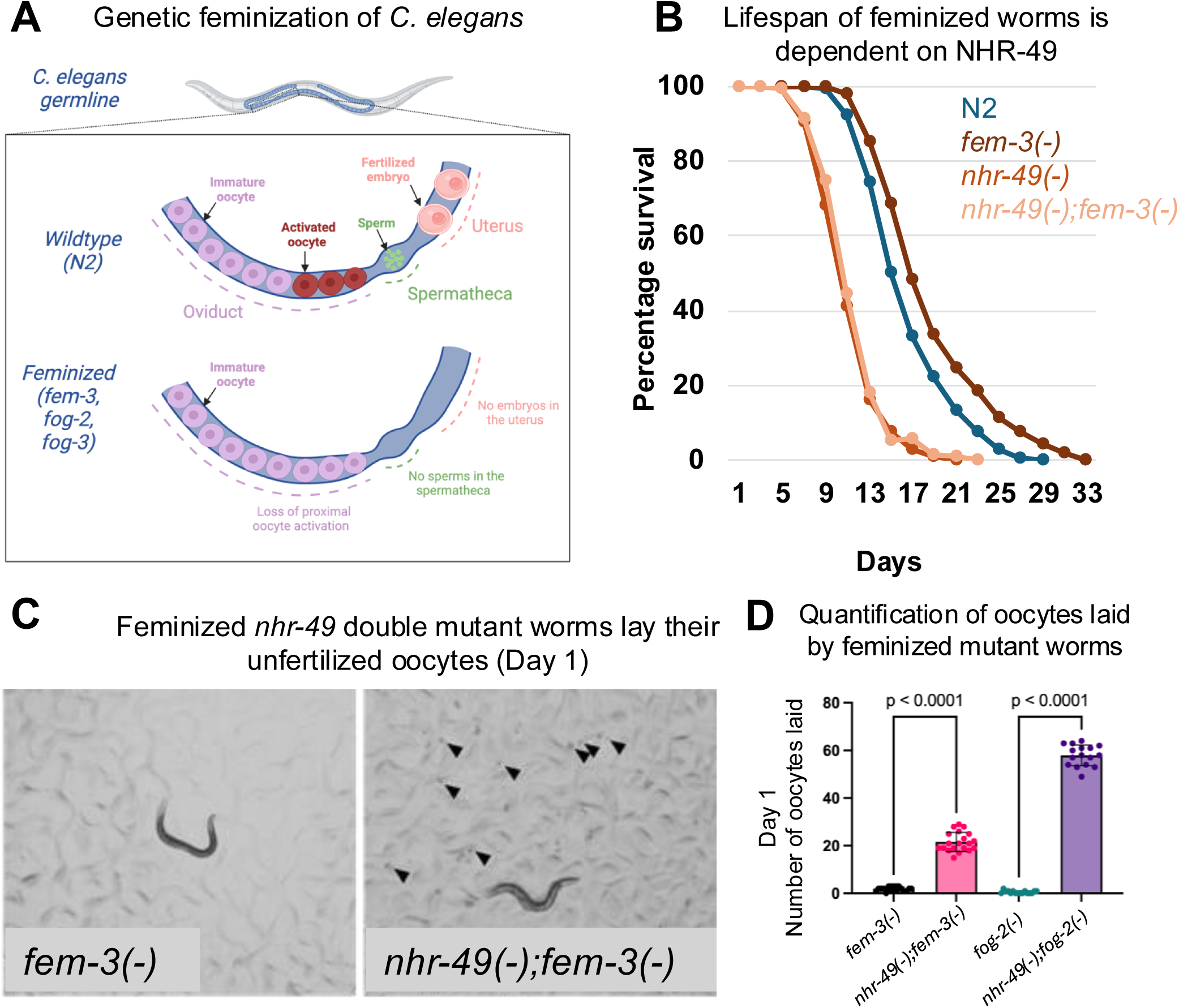
Loss of NHR-49 shortens lifespan and induces oocyte ovulation in feminized worms. **(A)** Schematic representation of feminization of *C. elegans* germline. **(B)** Survival curves for N2 (wild-type hermaphrodite), *nhr-49(gk405)* (hermaphrodite)*, fem-3(e1996) (*feminized*) and fem-3(e1996);nhr-49(gk405)*. Quantitative data and Kaplan-Meier statistics are provided in Table 1. **(C)** Representative images showing an absence of laid oocytes on plates with *fem-3(e1996)* mutants, and numerous oocytes (marked by arrowheads) were laid on plates with *fem-3(e1996);nhr-49(gk405)* double mutants on day 1 of adulthood. **(D)** Bar graphs showing the number of oocytes laid by *fem-3(e1996)* and *fem-3(e1996);nhr-49(gk405)* double mutants, and *fog-2*(*q71*) and *fog-2*(*q71*);*nhr-49(gk405)* on day 1 of adulthood. The number of oocytes laid were counted over a period of 24-hour period starting from the larval stage L4 (see Methods for details), (N >30) for each genotype, collected from two independent biological replicates. Statistical comparisons were performed using an unpaired t-test and are presented as mean ± SEM.

In *C. elegans*, the nuclear hormone receptor NHR-49—a putative functional homolog of mammalian PPARα and structural homolog of HNF4—is a central regulator of lipid metabolism and energy homeostasis^19^. Like all nuclear hormone receptors, NHR-49 is a DNA binding transcription factor that is regulated through ligand binding^20,21^. NHR-49 is known to regulate the expression of both lipid synthesis and lipid hydrolysis genes^22^. For example, NHR-49 transcriptionally regulates stearoyl-CoA desaturation genes *fat-6* and *fat-7,* which maintain membrane fluidity and polyunsaturated fatty acid synthesis^23^. It also activates *acs-2*, a gene involved in mitochondrial β-oxidation^19,23^. In addition to fat metabolism, NHR-49 is known to also regulate oxidative stress response^24^, hypoxia adaptation^25^, peroxisome proliferation^26^, pathogen avoidance^27^, innate immunity^28^, and longevity^29^. Interestingly, NHR-49 has been suggested to participate in regulating adult reproductive diapause, a state in which severely starved L4 worms undergo extensive germline attrition and the animal can survive for a period of several weeks ^30,31,32^, and the germline can regenerate when food become available again.

Using feminized mutant C. elegans as a model, we uncovered a previously unrecognized role for NHR-49 in regulating oocyte activation and ovulation in a cell non-autonomous manner. We observed that loss of *nhr-49* results in inappropriate oocyte activation and ovulation, and laying of unfertilized oocytes even in the absence of sperm signal. This inappropriate laying of unfertilized oocytes leads to excessive yolk and fat loss and drastically shortening of lifespan. Notably, when yolk transport to oocytes is blocked, thereby retaining yolk within the parent, fat storage is largely restored in nhr-49 mutant and their shortened lifespan is also partially rescued. Furthermore, *nhr-49* mutants failed to halt oocyte proliferation under nutrient-restricted conditions, suggesting a disruption in the organism’s ability to couple reproductive output to nutritional status. Together, our findings suggest that NHR-49 plays a broader role in coordinating energy metabolism and reproductive restraint, and in doing so, orchestrate the balance between parental resource management and reproductive success, with a major impact on longevity.

## Results

### NHR-49 is required for the longer lifespan and elevated fat storage of feminized *C. elegans*

Our lab recently reported that when not mated, feminized mutants exhibit significant lifespan extension and elevated fat storage compared to wild-type N2 (Wild-type) hermaphrodites at 25°C **(Fig. 1B)**^2^. While investigating the molecular factors required for the lifespan extension and fat metabolism of feminized worms, we identified NHR-49, a key metabolic regulator ^28,28,29,33,34^, as essential for the enhanced lifespan of feminized worms **(Fig. 1B)**. Loss of *nhr-49* drastically reduced the lifespan of feminized worms, where the double mutant *nhr-49(gk405);fem-3(e1996)* had lifespan similar to the short lifespan of the *nhr-49(gk405)* single mutant^33^.

While characterizing the *nhr-49(-);fem-3(-)* double mutant, we noticed a surprising and interesting phenotype, where these feminized mutants laid unfertilized oocytes during early adulthood (**Fig. 1C**). In *C. elegans*, oocyte maturation and ovulation are strictly dependent on the presence of sperm, and consequently, feminized mutants lacking sperm retain their oocytes within the oviduct^15^. Accordingly, we observed no oocytes laid on the plate for *fem-3(-)* mutants **(Fig. 1C and 1D)**^3^. When examining *nhr-49(-);fem-3(-)* double mutants, we observed that these worms laid their unfertilized oocytes even though they lack sperm **(Fig. 1 C and D)**. Similar observation was made with a different feminized mutant strain^35^ *fog-2(-)* and *nhr-49(-);fog-2(-)* **(Fig. 1D)**. NHR-49, and its homologs in other species, have not been implicated in regulating oocyte ovulation. To test whether NHR-49 uniquely regulates oocytes retention only in feminized mutants or whether it represents a more general role of NHR-49, we monitored *nhr-49(-)* mutant hermaphrodites. *C. elegans* hermaphrodites reach the end of their reproductive period around day 4 of adulthood at 20°C, when their sperm supply is depleted and the remaining oocytes are retained in the oviduct, mirroring the reproductive state of feminized mutants^36^. We first examined *nhr-49* single mutant hermaphrodites at this post-reproductive stage and observed that *nhr-49(-)* mutants also laid unfertilized oocytes when examined on day 6 of adulthood **(Fig. S1B)**, a phenotype similar that observed for *nhr-49(-);fem-3(-)*double mutants **(Fig. 1C)**. We additionally monitored egg laying of *nhr-49(-)* single mutant compared to wild-type hermaphrodites and observed that *nhr-49(-)* laid reduced number of fertilized embryos, but significantly higher number of unfertilized oocytes, throughout the reproductive period, as well as during the post-reproductive period (i.e. >120 hrs after reaching adulthood) **(Fig. S1B, S1D)**. This unexpected finding likely explains the previously reported lower brood size phenotype associated with the *nhr-49(-)* mutant^26^ (**Fig. S1A, C)**. Together, our results uncovered a new and fundamental role of NHR-49 in retention of unfertilized oocytes, with this function becoming particularly evident in feminized mutants where sperm is absent throughout development.

Different types of self-sterile *C. elegans* mutants, including feminized *fem-3(-)* mutants, accumulate excess fat^18^. Here, we found that the increased fat accumulation in *fem-3(-)* worms is dependent on NHR-49. We used Oil red O (ORO) staining^37^ to visualize storage lipid, and observed that both hermaphrodites and feminized mutant worms lacking *nhr-49* showed significantly lower ORO staining on day 6 adulthood **(Fig. 2 A and B)**. Interestingly, when comparing the ORO levels between day 1 and day 6 of adulthood in the different strains, wild type showed significant increase in stored fat as they progressed towards post-reproductive period, as previously published ^38–39^. *fem-3(-)* mutants showed no significant change during this timeframe. Hermaphrodites and feminized mutants lacking *nhr-49* showed drastic reduction in ORO from day 1 to day 6 **(Fig. 2 A and B)**, suggesting a depletion of stored fat in worms lacking *nhr-49*, which appears coincidental to the time-period where mutants lacking *nhr-4*9 are expelling their unfertilized oocytes **(Fig. S1)**. Adult *C. elegans* hermaphrodites are known to transfer lipid-bound yolk to their oocytes to support embryonic development^40^. Feminized worms have increased yolk accumulation due to a lack of fertilization and embryo formation, which can be visualized using VIT-2::GFP^41^, a reporter strain for one of the vitellogenin proteins. We observed that *nhr-49(-);fem-3(-)* double mutants, which lay their unfertilized oocytes, display reduced VIT-2::GFP signal by day 2 of adulthood, which parallel the reduced ORO in day 6 fem-3(-); *nhr-49(-)* double mutants **(Fig. S2A)**.

**Figure 2.**
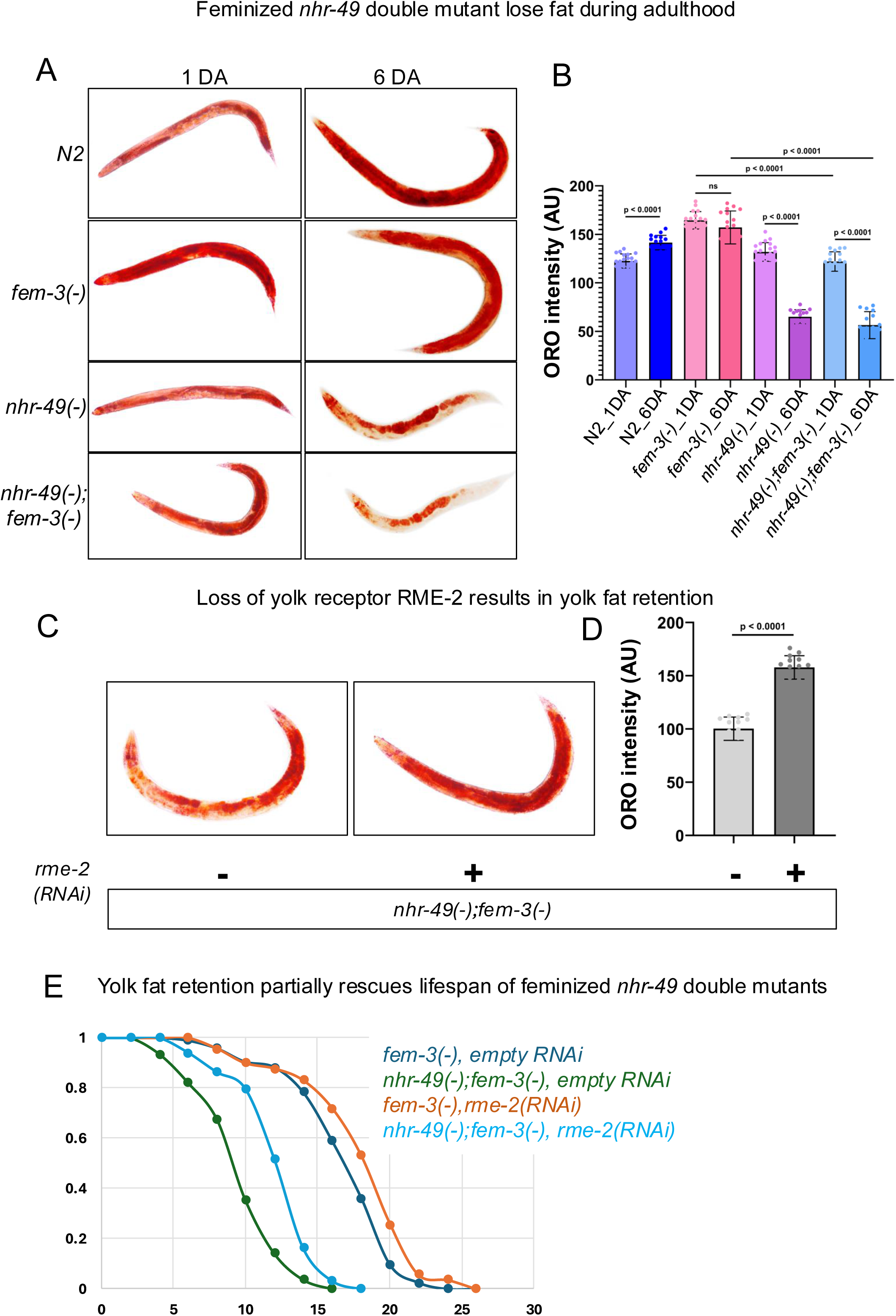
Lifespan shortening and fat loss in NHR-49 mutants are partially dependent on RME-2 mediated yolk transport into oocytes. **(A)** Representative images of Oil Red O staining in N2 wild-type (hermaphrodite), *fem-3(e1996)* (feminized)*, nhr-49(gk405)* (hermaphrodite), and *fem-3(e1996);nhr-49(gk405)* (feminized) worms on day 1 (1 DA) and day 6 (6 DA) of adulthood. **(B)** Oil red O (ORO) intensity in the indicated strains at day 1 and day 6 of adulthood. ORO intensity is shown as the mean intensity per-worm (N >30) for each genotype, collected from two independent biological replicates. Statistical comparisons were performed using one-way analysis of variance (ANOVA), followed by Tukey’s multiple comparisons test. **(C)** Loss of RME-2 restored fat accumulation, as shown by ORO staining. Representative images of ORO staining in *fem-3(-);nhr-49(-)* with and without *rme-2* RNAi on day 2 of adulthood. **(D)** Bar graphs showing ORO intensity in the indicated strains (day 1 adults). ORO intensity is shown as the mean intensity per-worm (N >30) for each genotype, collected from two independent biological replicates. Two-tailed p-values were computed using unpaired t-test. **(E)** Survival curves for *fem-3(e1996)*, and *fem-3(e1996);nhr-49(gk405)* double mutants on empty vector control and *rme-2* RNAi from one representative experiment. Quantitative data and Kaplan-Meier statistics are provided in Table 2.

Since both hermaphrodites^33^ and feminized mutants lacking *nhr-49(-)* are short-lived **(Fig. 1B)** and we found that they rapidly lose yolk by day 2 of adulthood **(Fig. S2)**, we hypothesized that attenuating yolk loss would help to preserve stored fat and lifespan. In *C. elegans*, yolk produced in the intestine is transported to oocytes via the endocytic receptor RME-2^42^. As expected, RNAi knockdown of *rme-2* resulted in yolk retention in *nhr-49(-);fem-3(-)* double mutants, observed as increased VIT-2::GFP reporter signal **(Fig. S2A and B)**. Yolk retention also largely restored fat storage, as indicated by ORO staining **(Fig. 2 C and D)**. Interestingly, knockdown of *rme-2* partially restored the lifespan of both *nhr-49(-);fem-3(-)* double mutants and *nhr-49(-)* single mutants **(Fig. 2E)**. Taken together, these findings indicated that laying of unfertilized oocytes is accompanied by shunting of yolk, which largely accounts for the rapid fat loss and partially accounts for the shortened lifespan of both hermaphrodites and feminized worms lacking *nhr-49(-)*.

### NHR-49 prevents oocyte activation and ovulation in the absence of sperm

We uncovered a new role of NHR-49 in preventing the laying of unfertilized oocytes, which appears to be coupled to the fat loss and shortened lifespan of *nhr-49(-)* mutants. To understand the molecular basis of inappropriate oocyte lay upon loss of *nhr-49*, we first characterized the activation status of the oocytes. In *C. elegans*, oocytes are naturally arrested in meiotic prophase I and require sperm-derived signals to resume meiotic progression, as part of oocyte maturation, ovulation, and subsequent fertilization. Doubly phosphorylated MPK-1 (dpMPK-1), a homolog of the ERK MAP kinase, is a well-established marker of oocyte activation^16^ **(Fig. 3A)**. We performed dpMPK-1 immunostaining of day-1 adults to examine the status of oocyte activation in hermaphrodites and feminized mutants with or without *nhr-49*. While fertile wild-type worms showed the expected dpMPK-1 signal in the oocytes proximal to the spermatheca, feminized mutants, both *fem-3(-)* and *fog-3(-)*^17^, did not **(Fig. 3A, B and S3A)**, indicative of a loss of oocyte activation in the absence of sperm. Loss of *nhr-49* did not substantially impact dpMPK-1 signal in reproductive hermaphrodites on day 1 of adulthood **(Fig. 3A and B)**. Interestingly, loss of *nhr-49* resulted in prominent dpMPK-1 signal in the proximal oocytes of feminized worms **(Fig. 3A, B, S3A, B)**, which was comparable to that in hermaphrodites, indicating inappropriate activation of the oocyte maturation pathway despite the absence of sperm. Similar inappropriate dpMPK-1 activation was also observed in post reproductive *nhr-49(-)* mutant hermaphrodites **(Fig. S3C, D)**.

**Figure 3.**
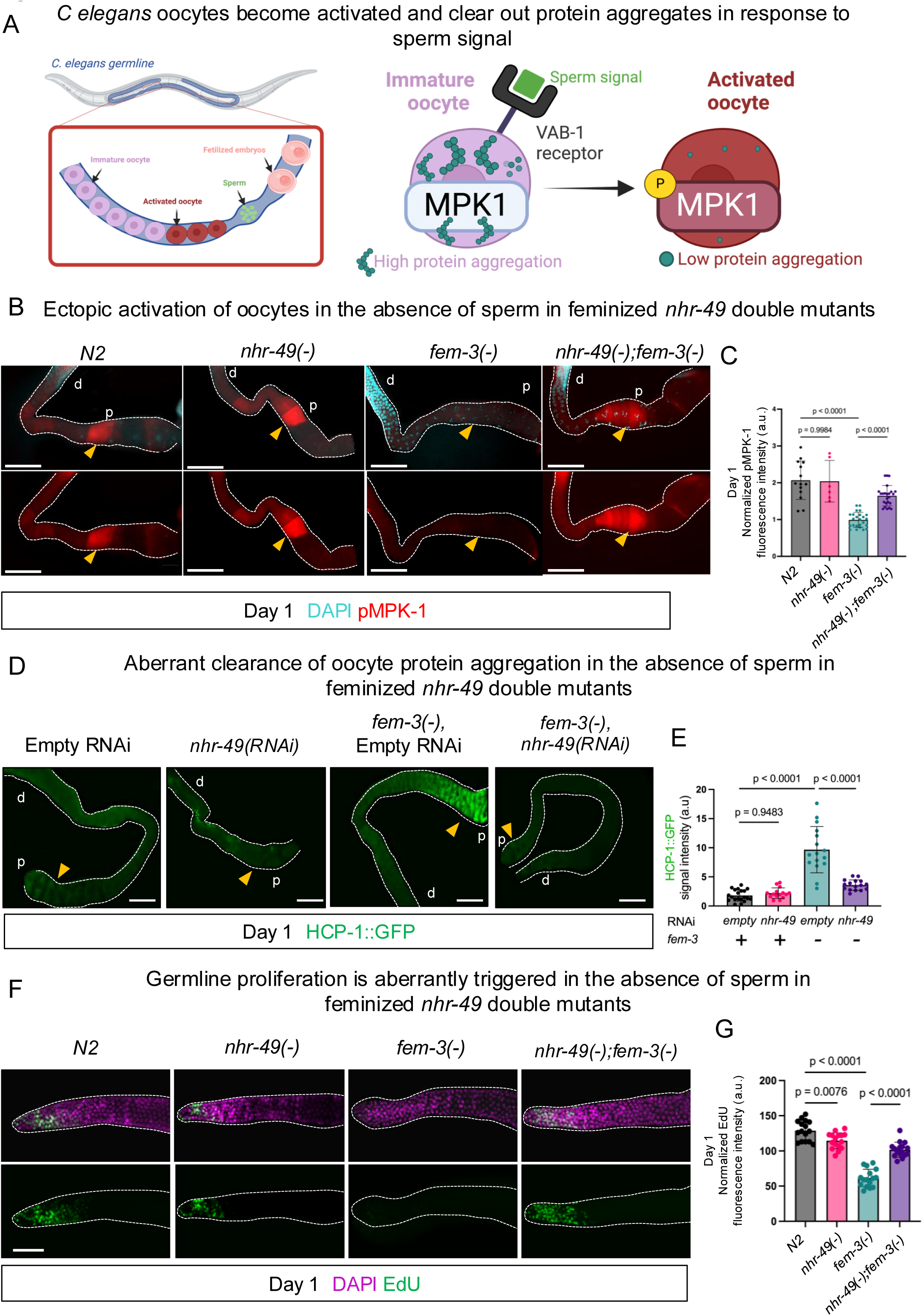
NHR-49 negatively regulates oocyte activation in feminized mutants. **(A)** Schematic diagram representing dpMPK-1 activation and protein aggregation states in the proximal region of the germline. **(B)** Representative images of dissected day-1 adult gonads immunostained with anti-dpMPK-1 antibody, which recognizes the di-phosphorylated, active form of MPK-1 (red). The position of the oocyte most proximal to the spermatheca in each image is marked by a yellow arrowhead. DAPI staining (blue) was used to indicate germ cell nuclei. **(C)** Bar graph showing dpMPK-1 signal intensity in the proximal oocytes of the indicated strains. Statistical comparisons were performed using one-way analysis of variance (ANOVA) followed by Tukey’s multiple comparisons test. More than 45 worms from 2 independent replicates per genotype were analyzed. **(D)** Representative images of dissected day-1 adult gonads from wild-type or *fem-3(-)* worms expressing HCP-1::GFP fusion protein, treated with control RNAi (EV) or *nhr-49* RNAi. The position of the oocyte most proximal to the spermatheca in each image is marked by a yellow arrowhead. ‘p’ and ‘d’ indicate distal or proximal region of the gonad respectively. **(E)** Quantification of GFP fluorescence intensity in wild-type and *fem-3(-)* worms expressing HCP-1::GFP fusion protein, treated with control RNAi (EV) and *nhr-49* RNAi. Statistical comparisons were performed using one-way analysis of variance (ANOVA) followed by Tukey’s multiple comparisons test. More than 45 worms from 2 independent replicates per genotype were analyzed. **(F**) Representative images of dissected day-1 adult gonads stained with EdU (green), marking proliferating nuclei, and counterstained with DAPI (purple) to visualize germ cell nuclei. Shown are gonads from N2, *nhr-49(gk405)*, *fem-3(e1996),* and *fem-3(e1996);nhr-49(gk405)* worms. (G) Bar graph showing the number of EdU positive nuclei in the indicated strains. Statistical comparisons were performed using one-way ANOVA followed by Tukey’s multiple comparisons test. More than 45 worms from two independent replicates per genotype were analyzed. All scale bars = 50μm.

Oocytes arrested in meiotic prophase I maintain minimal protein quality control activity, resulting in the accumulation of protein aggregates throughout the cytoplasm^5^. During oocyte maturation, proximal oocytes eliminate these protein aggregates to prepare for fertilization^43^ **(Fig. 3A)**. To further assess the activation status of the oocytes in mutants lacking *nhr-49*, we examined protein aggregate clearance in feminized strains with or without *nhr-49*, using the HCP-1::GFP reporter strain that has been previously used to detect protein aggregates in *C. elegans* oocytes^43^. In day-1 adults, as expected, *fem-3(-)* single mutants had significantly higher HCP-1::GFP signal **(Fig. 3B)** in their proximal oocytes compared to reproductive wild-type hermaphrodites, consistent with that the oocytes are not activated and accumulate protein aggregates. RNAi knockdown of *nhr-49* in feminized worms resulted in aberrant loss of HCP-1::GFP signal **(Fig. 3B)** in the proximal oocytes, further supporting inappropriate activation of the oocytes when *nhr-49* is lost.

In summary, in both post-reproductive hermaphrodites and feminized mutants, loss of *nhr-49* leads to misactivation of oocytes, as well as laying of unfertilized oocytes. We therefore conclude that the indiscriminate laying of oocytes in the absence of sperm upon loss of NHR-49 occurs through misactivation of oocytes and their subsequent ovulation.

In C. *elegans*, successful reproduction requires continuous oocyte production to replace those that are fertilized and laid. This process depends on two key processes: active proliferation of the mitotic stem cells in the distal region of the germline, and subsequent oogenesis in the meiotic region. Since unfertilized oocytes were laid in worms lacking *nhr-49*, we predicted that germline stem cell proliferation and/or oogenesis would be enhanced. We assessed mitotic proliferation by EdU labelling^44^, which marks nuclei undergoing DNA synthesis, and oogenesis by immunostaining for dpMPK-1. Feminized mutants normally produce only a fixed number of oocytes due to them being retained in the oviduct, and as expected, they showed reduced number of EdU+ nuclei in the mitotic region **(Fig. 3C)** and lower dpMPK-1 signal intensity in the meiotic region **(Fig. S4A)** compared to wild-type control. Examination of *nhr-49(-);fem-3(-)* double mutant worms revealed aberrant proliferation of the distal germline stem cells as evident by higher levels of EdU+ nuclei in the mitotic region and inappropriate oogenesis as indicated by increased dpMPK-1 staining in the meiotic region **(Fig. 3C and S4A)**. Consistently, post-reproductive *nhr-49* mutant hermaphrodites also showed a significant elevation of dpMPK-1 signal in the meiotic region compared to wild-type worms at the same age, suggestive of aberrant oogenesis **(Fig. S4B)**. Collectively, these data revealed that both proliferation of the mitotic stem cells and oogenesis are aberrantly activated upon loss of NHR-49, despite the absence of sperm. However, it is not clear whether aberrant activation of oocytes and subsequent laying of unfertilized oocytes triggered distal germline proliferation and oogenesis or *vice versa* in *nhr-49* mutants.

### NHR-49 is required to couple nutritional status to germline proliferation

NHR-49 is a well-established metabolic regulator, with particular importance in maintaining lipid homeostasis upon starvation ^23,45^. NHR-49 has also been suggested to participate in starvation-induced germline attrition, a process known as adult reproductive diapause^46^. We therefore wondered whether the role of NHR-49 in the germline, as detailed above, is also sensitive to nutritional status. We monitored germline proliferation in reproductive hermaphrodites using EdU^44^ labeling and, as expected, observed that germline proliferation was significantly reduced in response to starvation (24 hours) in wild-type worms. We found that *nhr-49(-)* single mutant showed reduced germline proliferation as evident by lower EdU+ nuclei in the distal region. Notably, we did not observe further reduction of EdU+ nuclei upon starvation in the *nhr-49(-)* mutants **(Fig. 4A),** suggesting that NHR-49 is required for coupling nutritional status to germline proliferation.

**Figure 4.**
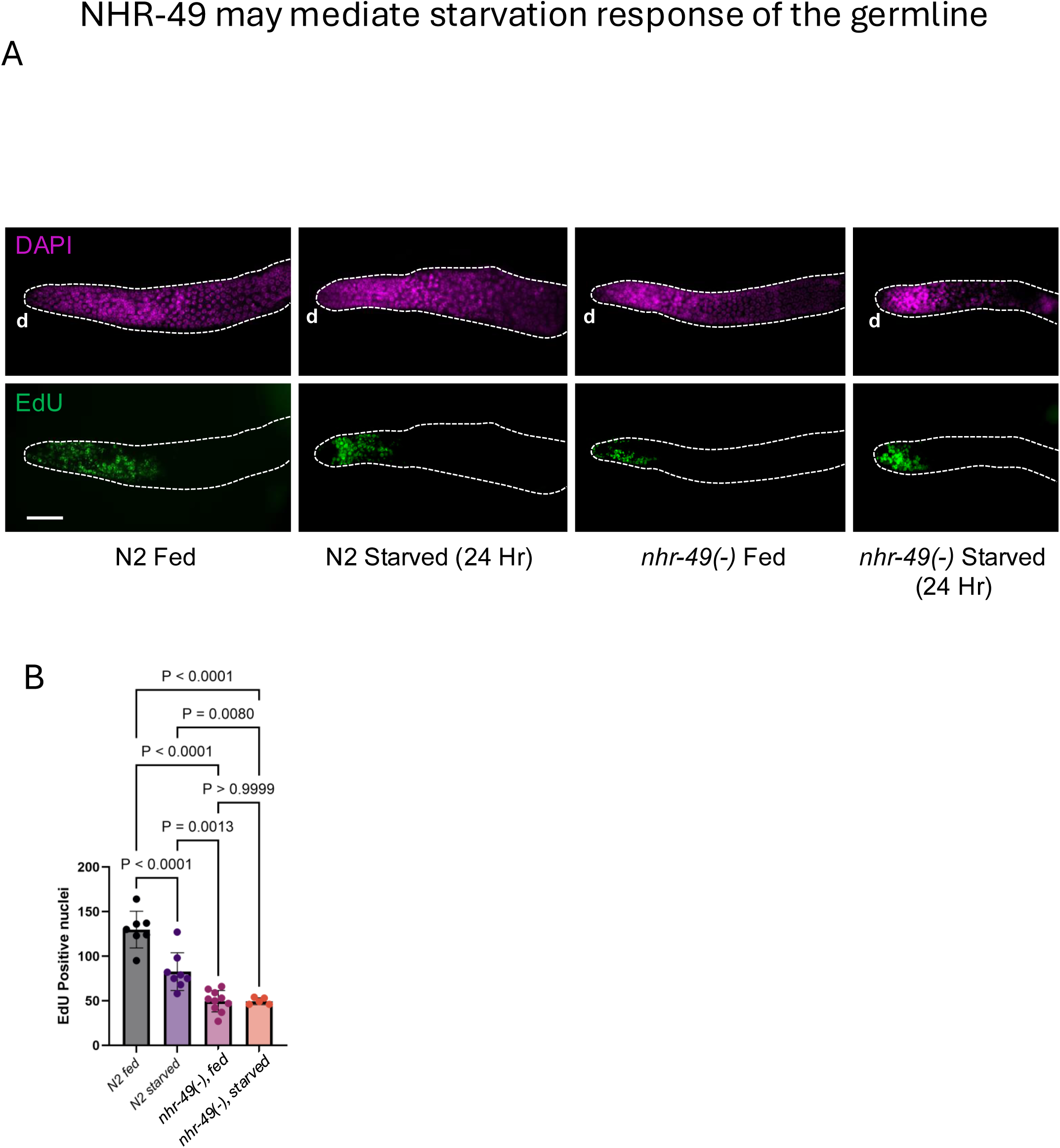
NHR-49 may mediate starvation response of the germline. **(A)** Representative images of EdU-stained dissected gonads from *N2* and *nhr-49(gk405)* worms under fed condition or after 24 hours of starvation started as late L4 stage and scored at day 1 of adulthood. Wildtype N2 germlines showed a significant decline in mitotic proliferation as indicated by EdU+ nuclei. *nhr-49(gk405)* mutant showed reduced EdU+ nuclei under fed condition but did not show further decrease in response to starvation. ‘d’ indicates distal region of the gonad. All scale bars = 50μm. **(B)** Bar graph showing the number of EdU+ nuclei in the indicated strains. Statistical comparisons were performed using one-way analysis of variance (ANOVA) followed by Tukey’s multiple comparisons test, based on more than 20 worms from two independent replicates per genotype.

### NHR-49 may regulate oocyte activation through the somatic gonadal sheath cells

To dissect the pathway through which NHR-49 regulates oocyte activation and ovulation, we examined its genetic interactions with known regulators of ovulation that act in distinct tissues. MSPs (Major Sperm Proteins) are sperm-derived cytoskeletal proteins that are also secreted as signaling molecules to promote oocyte maturation and sheath cell contraction^47^. *vab-1*, encoding an ephrin receptor expressed on oocytes, and *ceh-18*, a homeodomain transcription factor essential for sheath cell function, are both required for MSP-dependent oocyte activation. Given that sheath cells act as key MSP sensors, analogous to mammalian cumulus granulosa cells, these factors provide an entry point to determine the tissue of NHR-49 action. In feminized animals, RNAi knockdown of *vab-1* or *ceh-18* induces oocyte laying, where the effect of *vab-1* RNAi is modest, consistent with previous findings ^17^. Compared to *vab-1* RNAi, knockdown of *nhr-49* led to a significantly higher number of oocytes laid—nearly 6–10 times more than *vab-1* knockdown alone. Knockdown of *nhr-49* in *vab-1* mutants resulted in an oocyte-laying phenotype that was comparable to *nhr-49* RNAi alone **(Fig. 5A)**, suggesting a shared or converging pathway; However, given the mild effect of *vab-1* loss, it is also possible that an additive effect would be difficult to detect. Combining RNAi knockdown of *nhr-49* and *ceh-18* mutation also did not show additive effect, again suggesting that *nhr-49* functions in the same pathway as *ceh-18* **(Fig. 5B)**. These results suggested that NHR-49 might act in both the oocytes and the sheath cells to regulate oocyte ovulation.

**Figure 5.**
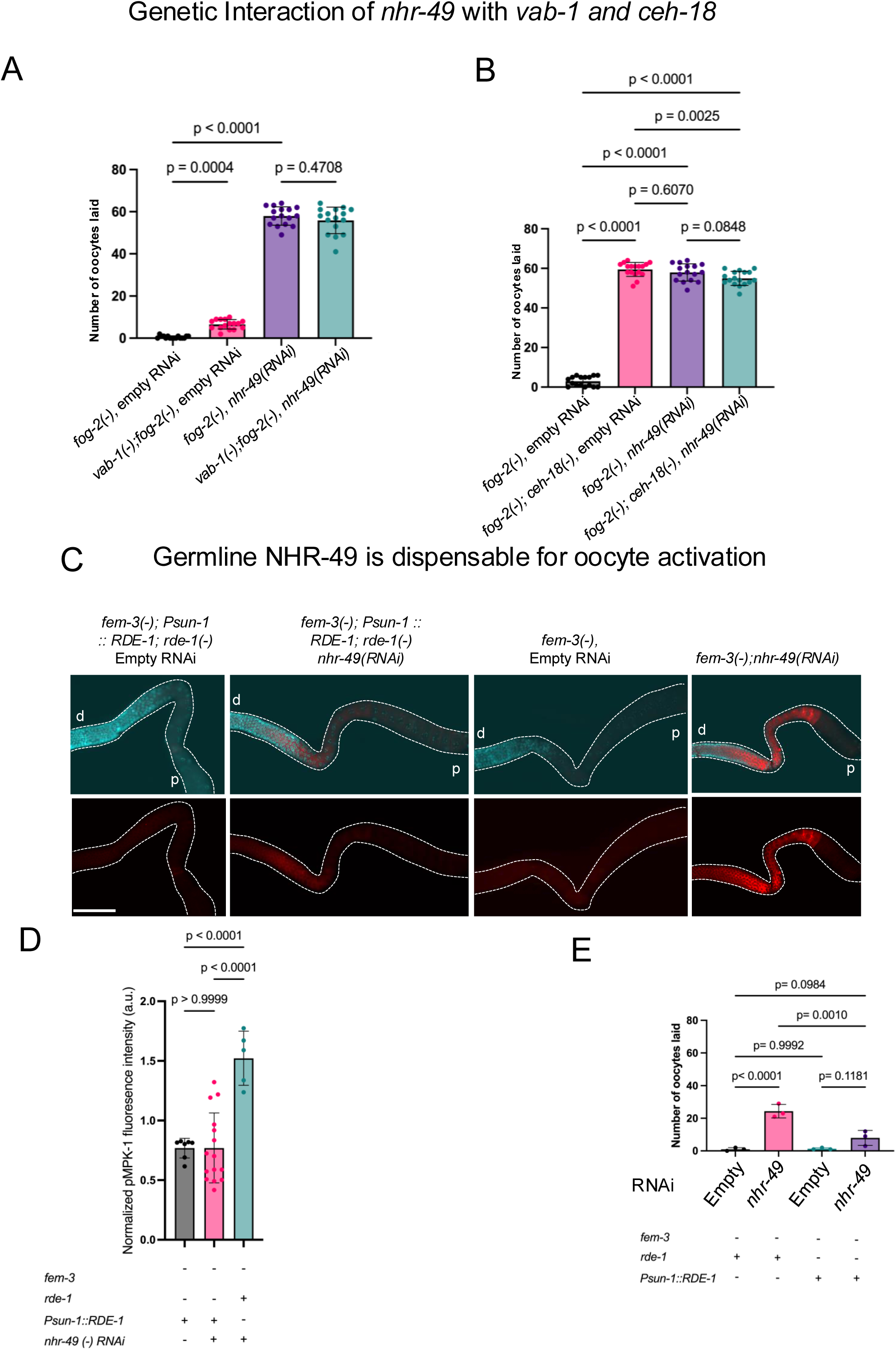
NHR-49 primarily regulates oocyte activation through somatic tissues. **(A)** Bar graph showing the number of unfertilized oocytes laid by *fog-2(q71), vab-1(dx31);fog-2(q71)* mutants with and without *nhr-49* RNAi. **(B)** Bar graph showing the number of unfertilized oocytes laid by *fog-2(q71), fog-2(q71);ceh-18(mg57)* with and without *nhr-49* RNAi. Comparisons were statistically tested by one-way analysis of variance (ANOVA) followed by multiple comparisons test with Tukey’s correction for >45 worms from 2 replicates for each genotype. **(C)** Representative images of dpMPK-1 immunostained dissected gonads from day-1 feminized mutants with either whole-body RNAi knockdown of *nhr-49* (right column) or germline-specific RNAi knockdown (second column) using *Psun-1::RDE-1* strain (see Methods for details of the germline-specific RNAi strain). dpMPK-1 signal (red) was observed in feminized mutants upon whole-body knockdown of *nhr-49,* but not when *nhr-49* was specifically knocked down in the germline. **(D)** Bar graph showing dpMPK-1 signal intensity in the proximal oocytes of the indicated strains. *rde-1* ‘+’ indicates a strain background that can undergo whole-body RNAi; *Psun-1::RDE-1* ‘+’ indicates a strain background that can undergo germline-specific RNAi. Comparisons were statistically tested by one-way analysis of variance (ANOVA) followed by Tukey’s multiple comparisons test. More than 25 worms from two independent replicates per genotype were analyzed. **(E)** Bar graph showing the number of unfertilized oocytes laid in strains undergoing whole-body ( *rde-1* ‘+’) or germline specific (*Psun-1::RDE-1* ‘+’) RNAi knockdown of *nhr-49*. Comparisons were statistically tested by one-way analysis of variance (ANOVA) followed by Tukey’s multiple comparisons test. More than 25 worms from two independent replicates per genotype were analyzed. All scale bars = 50μm.

To further test whether NHR-49 acts cell-autonomously in oocytes or from somatic tissues to regulate oocyte ovulation, we used a germline-specific RNAi approach, where *rde-1* null mutant is complemented with functional RDE-1 only in the germline (*rde-1(-)*; *Psun-1*::RDE-1)^48^. Germline-specific knockdown of *nhr-49* in feminized worms failed to recapitulate the oocyte-laying phenotype and dpMPK-1 misregulation observed with whole-body *nhr-49* loss **(Fig. 5C–E)**. These results indicate that NHR-49 is largely dispensable in the germline and instead acts through somatic tissues—likely the sheath cells—to control oocyte activation.

### Genome-wide analyses identify metabolic and stress-related transcriptional targets of NHR-49

To further investigate the transcriptional outputs of NHR-49, we performed CUT&RUN, a chromatin profiling method that maps protein-DNA interactions, in day 1 adult wild-type animals using the CRISPR-engineered *nhr-49(hj293)* strain, which harbors a GFP::TEV::3xHA tag at the C - terminus of the endogenous *nhr-49* locus ^26^. This profiling revealed 2,227 high-confidence NHR-49 binding sites, with the majority of peaks localized to gene promoters and transcription start sites **(Fig. S8A–B, Table 3)** consistent with a role of NHR-49 in transcriptional regulation. Gene set enrichment analysis ^49^ of NHR-49-bound genes highlighted key categories related to lipid metabolism, mitochondrial fatty acid oxidation, and detoxification pathways **(Fig. S8C–D)**.

To assess the functional impact of NHR-49 binding, we performed RNA-seq on *nhr-49(-)* mutants and wild-type controls at the day 1 of adulthood. Differential expression analysis revealed significant up- and downregulation of genes associated with lipid metabolism, innate immunity, and extracellular matrix remodeling **(Fig. S7A–B, Table 4)**. As an additional quality control, we compared our *nhr-49(-)* RNA-seq data to previously published transcriptomic profiles of *nhr-49* mutants. We found that the three data sets showed significant overlap, confirming a shared core set of NHR-49–dependent gene expression changes across studies **(Fig. S7E–F)**. However, each dataset also displays distinct subsets of regulated genes, suggesting that chronic loss of *nhr-49* function can lead to compensatory or context-specific transcriptional adaptations.

Integrating CUT&RUN and RNA-seq data revealed 27 genes that were both bound by NHR-49 and differentially expressed in *nhr-49* mutants **(Fig. S7C, Table 5)**, suggesting they represent direct transcriptional targets. Gene set enrichment analysis indicated that the upregulated genes are enriched for lipid biosynthesis and pathogen stress responses, and the downregulated genes are enriched for mitochondrial function.

### NHR-49 represses *gsa-1* expression to prevent inappropriate oocyte activation

Among the NHR-49-bound targets, we identified *gsa-1*, which encodes a Gαs subunit known to act in the gonadal sheath cells to mediate MSP signaling and regulate oocyte ovulation ^50^. NHR-49 binding peaks were detected at the promoter and second intron of *gsa-1* **(Fig. 6C)**. Interestingly, RNA-seq indicated that *gsa-1* RNA expression was modestly elevated in *nhr-49* mutant **(Fig. 6A)**, albeit not reaching significance.

**Figure 6.**
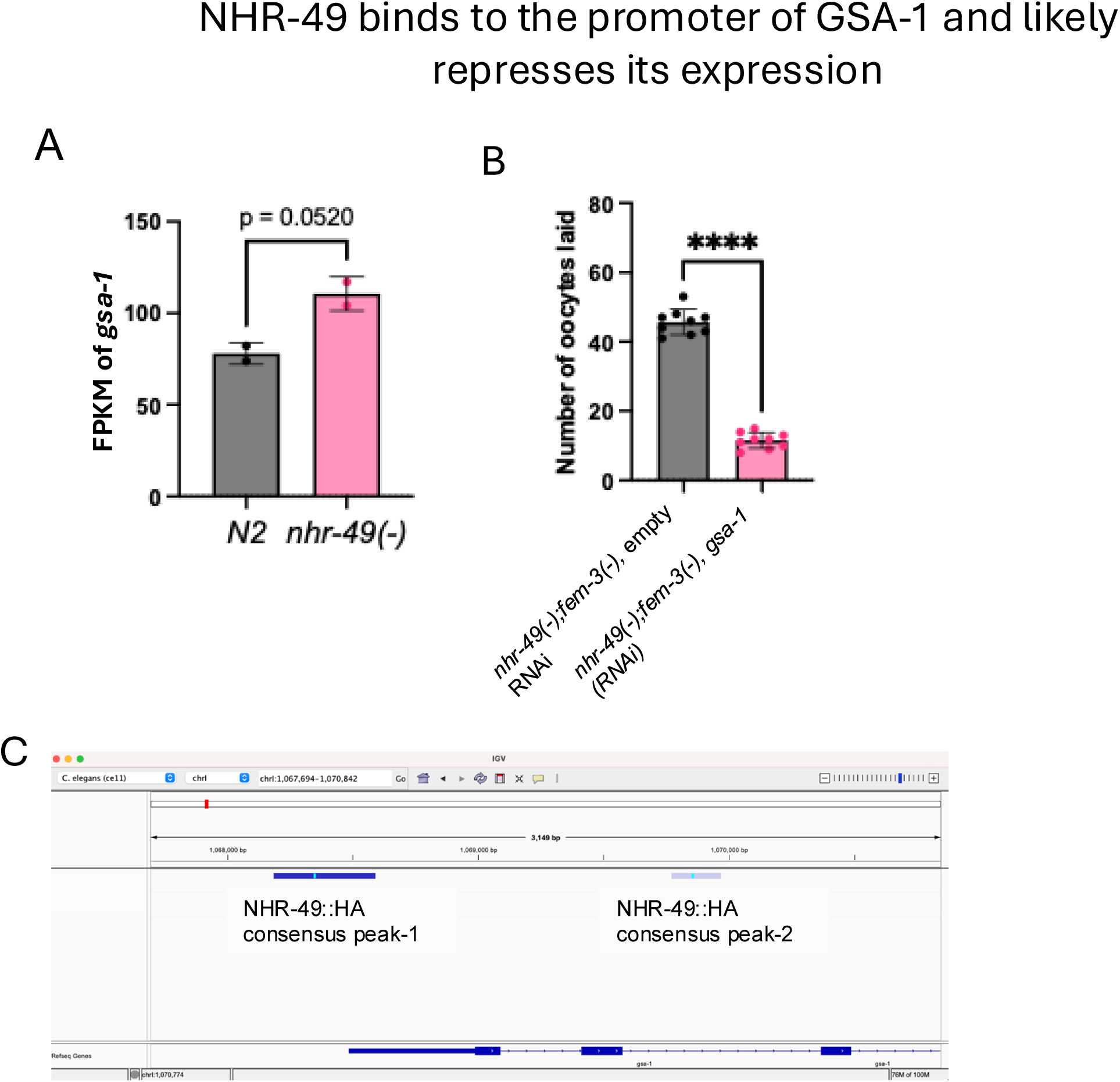
NHR-49 may regulate oocyte activation by repressing sheath cell-mediated sperm sensing. **(A)** Bar graph showing a trend of elevated *gsa-1* mRNA expression in *nhr-49(gk405)* worms compared to *N2* (wild-type) based on RNAseq results. Two-tailed p-values were computed using an unpaired t-test. N >30 2 independent biological replicates **(B)** Bar graph showing the number of unfertilized oocytes laid by *fem-3(e1996);nhr-49(gk405)* double mutants treated with control (EV) and *gsa-1* RNAi. Two-tailed p-values were computed using an unpaired t-test. N >30 2 independent biological replicates **(C)** Screenshot from Integrative Genomics Viewer (IGV) showing NHR-49-HA binding at the promoter and intragenic regions of *gsa-1*, as detected by CUT&RUN chromatin profiling. Darker blue at promoter region indicates stronger signal with signalValue: 2.26408 pValue: 82.6164 qValue: 79.556 and lighter blue signal at the intron region suggests comparatively weaker signal with a signalValue: 1.49912 pValue: 5.58139 qValue: 4.01701. Fluorescent blue signal in the middle of the peak indicates peak summit.

Importantly, functional experiments showed that RNAi knockdown of *gsa-1* suppressed the excessive oocyte laying phenotype of *nhr-49(-);fem-3(-)* animals **(Fig. 6B)**, indicating that elevated gsa-1 expression likely contributes to the aberrant laying of unfertilized oocytes. Taken together, the data suggested that NHR-49 acts, at least in part, through repressing *gsa-1* expression to constrain inappropriate oocyte activation.

## Discussion

Our findings uncover a previously unrecognized role for the nuclear hormone receptor NHR-49 in regulating oocyte activation and ovulation in *C. elegans*. This reproductive role appears to be mediated through the transcriptional repression of *gsa-1*, a Gαs protein that promotes oocyte activation, and likely occurs via the gonadal sheath cells rather than the germline itself. We find that inappropriate oocyte activation and laying of unfertilized oocytes lead to premature yolk loss and depletion of parental resources.

Suppressing yolk loss can partially restore lifespan, linking the reproductive phenotype to diminished somatic maintenance, and offers a partial explanation of the previously described lifespan shortening phenotype of the *nhr-49* mutant. While NHR-49 has been extensively characterized as a regulator of lipid metabolism and lifespan, our data add a new dimension to the understanding of how metabolic regulators like NHR-49 influence longevity—not only by altering fat metabolism, but by controlling the timing and scale of reproductive investment.

Our data suggest NHR-49 is required to suppress germline proliferation in adult worms subjected to acute nutrient deprivation. We found that *nhr-49* mutants fail to reduce mitotic activity^4^ in the distal germline following short-term starvation, suggesting that NHR-49 mediates nutrient-responsive germline restraint. NHR-49 is also known to be required for adult reproductive diapause (ARD)^30^—a distinct starvation-induced state in L3/L4 animals characterized by drastic germline shrinkage. While the mechanisms underlying these two responses may differ, the shared requirement for NHR-49 raises the possibility that it acts as a conserved metabolic regulator coordinating reproductive quiescence in response to energetic stress. Given that specific dietary and microbiota-derived metabolites can activate NHR-49^40^, these findings support a broader model in which NHR-49 functions as a metabolic sensor linking environmental inputs to germline decisions.

Our CUT&RUN profiling revealed that NHR-49 binds near the loci of over 100 other nuclear hormone receptors, suggesting that it may serve as a broader transcriptional regulator of the NHR network itself. This opens the possibility that NHR-49 tunes hormonal signaling pathways across tissues in response to nutrient state. Future CUT&RUN studies in fed versus starved animals could determine whether NHR-49 differentially engages or recruits other NHRs depending on metabolic conditions. Additionally, future studies employing acute depletion system will likely reveal the immediate, direct transcriptional responses to NHR-49 loss from secondary effects that accumulate in constitutive mutants.

NHR-49 regulates *gsa-1*, a Gαs protein that functions in the gonadal sheath cells to promote oocyte activation and ovulation. Germline-specific RNAi suggests that NHR-49 does not act through the germline to regulate laying of unfertilized oocytes. Based on the known site of action of GSA-1 in the gonadal sheath cells, the observation that *gsa-1* expression appears elevated in *nhr-49* mutants, and the binding of NHR-49 to the promoter and intron regions of *gsa-1*, we suggest that NHR-49 constrain oocyte activation and ovulation in the absence of sperm by repressing *gsa-1* transcription in the sheath cells. Future tissue-specific studies will be important to confirm this proposed site of action.

Together, these findings position NHR-49 as a metabolic gatekeeper that couples environmental and internal nutrient status to reproductive restraint. By limiting oocyte activation and germline proliferation when resources or sperm are lacking, NHR-49 helps align reproductive effort with metabolic capacity, promoting long-term survival and maternal resource preservation.

## Material and Methods

### *C. elegans* culture methods and strains

*C. elegans* strains were maintained using standard protocols. Unless otherwise stated, all strains were grown on nematode growth medium (NGM) plates seeded with E. coli OP50-1 as a food source. The following strains were obtained from the Caenorhabditis Genetics Center (CGC): N2, DG2189 [*fog-3(q443) I/hT2 [bli-4(e937) let-?(q782) qIs48] (I;III); tnIs13 [pie-1p::vab-1::GFP + unc-119(+)] ltIs44 [pie-1p::mCherry::PH(PLC1delta1) + unc-119(+)*]; DG2190 [*fog-3(q443) I/hT2 [bli-4(e937) let-?(q782) qIs48] (I;III); tnIs13 [pie-1p::vab-1::GFP + unc-119(+)]V*]; VC470 [*nhr-49 (gk405)*]; DCL569 [*mkcSi13[sun-1p::rde-1::sun-1 3’UTR + unc-119(+)] II; rde-1(mkc36)V*]. These strains were previously generated in our laboratory, combining strains obtained from the CGC: IU613 [*fem-3(e1996)/nT1 [qIs51] (IV;V)]; IU614 [fem-3(e1996)/nT1 [qIs51] (IV;V)*]

### Lifespan Analysis

Lifespan assays were performed as previously described^18^. 10-15 gravid worms were placed on plates containing the desired bacterial food to obtain approximately 150-200 embryos. Upon reaching day one of adulthood, 30 worms were transferred to each of three RNAi plates and monitored every other day. Worms were gently transferred to fresh plates every other day to prevent starvation and mixing of generations. Care was taken to avoid transferring larvae. Lack of movement in response to gentle touch at the nose was used to confirm death. Worms that died due to rupturing, bursting, or leaving the plate were considered censored. The lifespan data was analyzed using the OASIS online survival analysis tool^78^ available at https://sbi.postech.ac.kr/oasis2/, and log-rank tests were used to determine statistical significance between groups. All lifespan assays were performed at least twice.

### RNAi Treatment

The RNAi procedure was carried out as previously described^18^. RNAi was conducted by feeding the worms with HT115 bacteria that expressed double-stranded RNA of the target gene. The RNAi clones were obtained from RNAi feeding libraries created by Julie Ahringer and Marc Vidal’s laboratories^76,77^.

### pMPK-1 Immunostaining

Approximately 100 worms were grown on OP50-1 seeded NGM plates, then washed off using PBST (PBS + 0.1% Tween 20) and collected in a 1.5 ml Eppendorf tube. The worms were spun for 15 seconds at 2000 rpm, the supernatant was removed, and the pellet was washed with 1X PBS twice, using 1 ml each time. Approximately 50 worms in 100 µl of PBS buffer were then transferred using a cut 200 µl micropipette tip onto a border-painted glass slide. To anesthetize the worms, 5 µl of 5 mM levamisole was added. Using 25- or 26-gauge needles, a cut was made near the pharyngeal bulb of each worm to release gonads, and all animals were dissected within 5–6 minutes. Dissected gonads were collected into a 1.5 ml tube using PBS-Tween–coated, cut-end 200 µl tips. Excess liquid was removed, and 200 µl of 4% formaldehyde fixative (prepared by mixing 100 µl 10X phosphate buffer, 100 µl 37% formaldehyde, and 800 µl ddH₂O to make a total volume of 1 ml fixative) was added. Samples were incubated at room temperature for 20-30 minutes. Fixative was then removed, and worms were incubated in 1 ml of chilled methanol (–20 °C) for 5 minutes. After removing methanol, worms were washed three times with PBST (5 minutes each). Excess liquid was removed, and worms were resuspended in 200 µl blocking buffer (0.1% Triton X-100, 0.1% Tween-20 and 2% BSA (Bovine Serum Albumin) in 1X PBS) and incubated for 1–2 hours. Primary antibody diluted 1:400 in Blocking buffer (50 µl per sample) was added and incubated overnight at 4 °C with rocking. Samples were then washed three times with blocking buffer for 10 minutes each. Secondary antibody (1:200 dilution in blocking buffer, 50 µl per sample) along with DAPI (final concentration 5 µg/ml) was added, and samples were incubated for 2 hours at room temperature in the dark with rocking. Following three washes with blocking buffer, worms were resuspended in 10 µl of Vectashield mounting medium. Worms were mounted on a 2% agarose pad prepared on a glass slide and covered with a coverslip for microscopy.

### EdU (5-ethynyl-2’-deoxyuridine) Staining

EdU staining was performed as previously described, with modifications^82^, using the Click-iT® EdU Imaging Kit (ThermoFisher Cat. No. C10337). For EdU incorporation, 50-100 worms were washed and collected from plates using M9 buffer containing 0.1% Triton X-100 (M9T) and washed twice with 1 mL M9T. 50 µL of M9T containing the worms were transferred into a fresh 1.5 mL microcentrifuge tube using a cut tip, followed by the addition of 50 µL of 10 mM EdU. The worms were gently mixed and incubated on a shaker for 40 minutes at room temperature (22–25°C). Worms were washed twice with 1 mL M9T. Next, 30 µL of EBT (prepared by mixing 110 µL 10X egg buffer, 10 µL 10% Tween-20, 6.5 µL 10% tetramisole, and 850 µL distilled water to a final volume of 976.5 µL) was pipetted onto a slide, and approximately 50 worms were transferred into the EBT drop. The worms were anesthetized for 10-15 seconds, and their heads were cut to extrude the gonads. The gonads were collected into a 1.5 mL microcentrifuge tube, fixed with 4% formaldehyde in egg buffer for 20-30 minutes, and washed twice with 1 mL of 3% BSA in PBS. For permeabilization, the gonads were incubated in 1 mL of 1% Triton X-100 in PBS at room temperature for 20 minutes. After permeabilization, the buffer was removed, and the samples were washed twice with 1 mL of 3% BSA in PBS. Subsequently, 0.5 mL of the Click-iT® reaction cocktail (as per the kit instructions) was added, and the worms were incubated at room temperature for 30 minutes on a shaker and covered to protect from light. The reaction cocktail was removed, and the samples were washed twice with 1 mL of 3% BSA in PBS. For the EdU chase experiment, after the 10 mM EdU incorporation step, worms were transferred to the feeding plates for 24 hours, and the subsequent steps were performed as described above. Finally, 10 µL of Vectashield was added, and the worms were mounted on a 2% agarose pad prepared on a glass slide for imaging. Images were acquired using a Keyence BZ-X810 microscope.

### Fluorescence quantification and imaging

For imaging, 10-20 worms were immobilized on a 2% agarose pad prepared on a glass slide in 10 µL of 5 mM levamisole in PBS. Images were acquired using the appropriate fluorescence channel on Leica DM5000B compound microscope or a Zeiss LSM 710 or LSM 880 inverted confocal microscope. Images were acquired as Z-stacks, and image tiling was used to scan at 63X magnification under oil immersion. For data processing, ZEN software was used for image stitching and FIJI and was used for image analysis and processing. To generate representative images, Adobe Photoshop was used.

### RNA isolation and library preparation for RNA-seq

RNA isolation and RNA-seq library preparation were performed as previously described ^18^. For the collection of whole worm samples, approximately 200 worms were collected in TRI reagent (MRC, Cat. No. TR 118). For tissue-specific analysis, germlines were surgically isolated from 100-150 animals using fine-gauge needle dissection, with 80-100 intact germlines collected per sample in TRI reagent. Two to three biological replicates were collected for each genotype. RNA-seq libraries were prepared using the QuantSeq 3‘ mRNA-Seq Library Prep Kit for Illumina (FWD) (Lexogen), and sequencing was performed by the Biotechnology Resource Center (BRC) of the Cornell Institute of Biotechnology.

### RNA-seq data analysis

RNA-seq data analysis was performed as previously described ^18^ with minor modifications. To obtain the total read count of RNA-seq data were processed using Linux on the BioHPC server at Cornell University. Raw sequencing data underwent quality control and preprocessing using Trim Galore (v0.6.5), which incorporates Cutadapt^51^ (v3.4) and FastQC (v0.11.8). Parameters were set to retain reads with quality scores ≥20 (-q 20) while performing FastQC quality assessment. Preprocessed reads were mapped to the *C. elegans* reference genome (ce11/WBcel235) using STAR aligner (v2.7.9a)^52^, with parameters configured for strict alignment: two computing threads (--runThreadN 2), gene count quantification (--quantMode GeneCounts), unique mapping only (--outFilterMultiMapNmax 1), a maximum of two mismatches (-- outFilterMismatchNmax 2), and coordinate-sorted BAM files (--outSAMtype BAM SortedByCoordinate). For 3’ RNA-seq data analysis, the middle column of counts from the STAR-generated tab-delimited files was used to construct expression matrices for comparative analysis. Matrices were organized either by genotype (comparing different mutant strains under identical conditions) or by treatment (comparing identical genotypes under different RNAi conditions). Differential expression analysis was performed in R (RStudio) using DESeq2 (v1.34.0)^53^. To improve statistical power, genes with fewer than 10 reads across fewer than 2 samples were excluded. Principal component analysis was conducted on variance-stabilizing transformed data (using the vst function), visualized using the plotPCA function with default parameters, and further refined using ggplot2 (v3.3.6). Statistically significant differentially expressed genes were identified using the standard DESeq2 workflow with an adjusted p-value threshold of 0.05. For genes of interest, normalized counts were visualized using DESeq2’s plotCounts function. Gene set comparisons between genotypes or between RNA-seq and CUT&RUN datasets were performed using DeepVenn (https://www.deepvenn.com/)^54^ online software, with statistical significance determined by Fisher’s exact test. Gene set enrichment analysis was conducted using WormCat 2.0 (www.wormcat.com)^49^, which also employs Fisher’s test with FDR correction. For heatmap generation, log_2_ fold-change values were extracted from DESeq2 results tables without filtering out low-expression genes. Heatmaps were created using the ComplexHeatmap package (v2.10.0) with row clustering enabled (cluster_rows = TRUE), column clustering disabled (cluster_columns = FALSE), and row names hidden (show_row_names = FALSE). For clustered analyses, heatmaps were partitioned (split = 3), and genes from each cluster were extracted using the row_order function.

### CUT&RUN for NHR-49 binding sites in the genome

CUT&RUN experiments were performed using our previously published protocol^55,56^. A sufficient number of embryos were collected from gravid adult worms grown on 10-cm NGM plates seeded with OP50-1. For each CUT&RUN reaction, 3000 embryos were plated onto 15-cm NGM plates supplemented with 1 mL of 25× concentrated OP50-1 culture (grown overnight). Sample preparation was scaled to ensure sufficient worm numbers (3000 animals per reaction) for each experimental condition. Plates were incubated at 25°C for approximately 48-52 hours until populations reached the predominantly young adult stage. At this developmental stage, worms were harvested by washing plates with PBS buffers, and CUT&RUN experiments were performed according to a detailed protocol previously published^55^ which follows standard CUT&RUN methodology with minor modifications specific to *C. elegans* samples. Sequencing libraries were constructed using the NEBNext Ultra II DNA Library Prep Kit for Illumina (NEB, catalog numbers E7645S and E7335S). Library amplification was performed with 14 PCR cycles according to the manufacturer’s guidelines, with slight procedural adjustments as described in the published protocol^57^. The resulting libraries underwent paired-end sequencing (2 × 32 bp) on an Illumina NextSeq 500 platform, performed by the Biotechnology Resource Center (BRC) of the Cornell Institute of Biotechnology. All CUT&RUN experiments were conducted with at least two independent biological replicates.

### CUT&RUN Analysis

CUT&RUN analysis was performed as previously described^55,58,59,60,57^.

## Supporting information

Zipped_Supplemental_Files

## Acknowledgments

We would like to thank personnel at Cornell Genomics Center, Microscopy core and statistical consulting, and the Caenorhabditis Genetics Center (CGC) for all the worm strains used in this study. We thank B. Sanketi, A. Ren, N. Kurpios, R. Sardana, K. Liu, D. Lin, G. Hollopeter members of the Cornell worm club, P. Cohen and members of the Cornell Reproductive Sciences Center, and the members of the Lee lab for their valuable discussion and feedback.

## Author contributions

Conceptualization, S.G., A.C., S.S.L.; Methodology, S.G., A.C., S.S.L.; Investigation, S.G., A.C.; Investigation: assistance with selected experiments, T.C., J.O., R.R., Data analysis, S.G., A.C.; Data curation, S.G., A.C.; visualization, S.G., A.C., S.S.L.; funding acquisition, S.S.L.; supervision, S.S.L.; writing – original draft, S.G.; and writing – review and editing, A.C., S.S.L.

## Materials and correspondence

Requests for materials may be directed to (SSL).

## Supplementary figure legends

**Figure S1.**
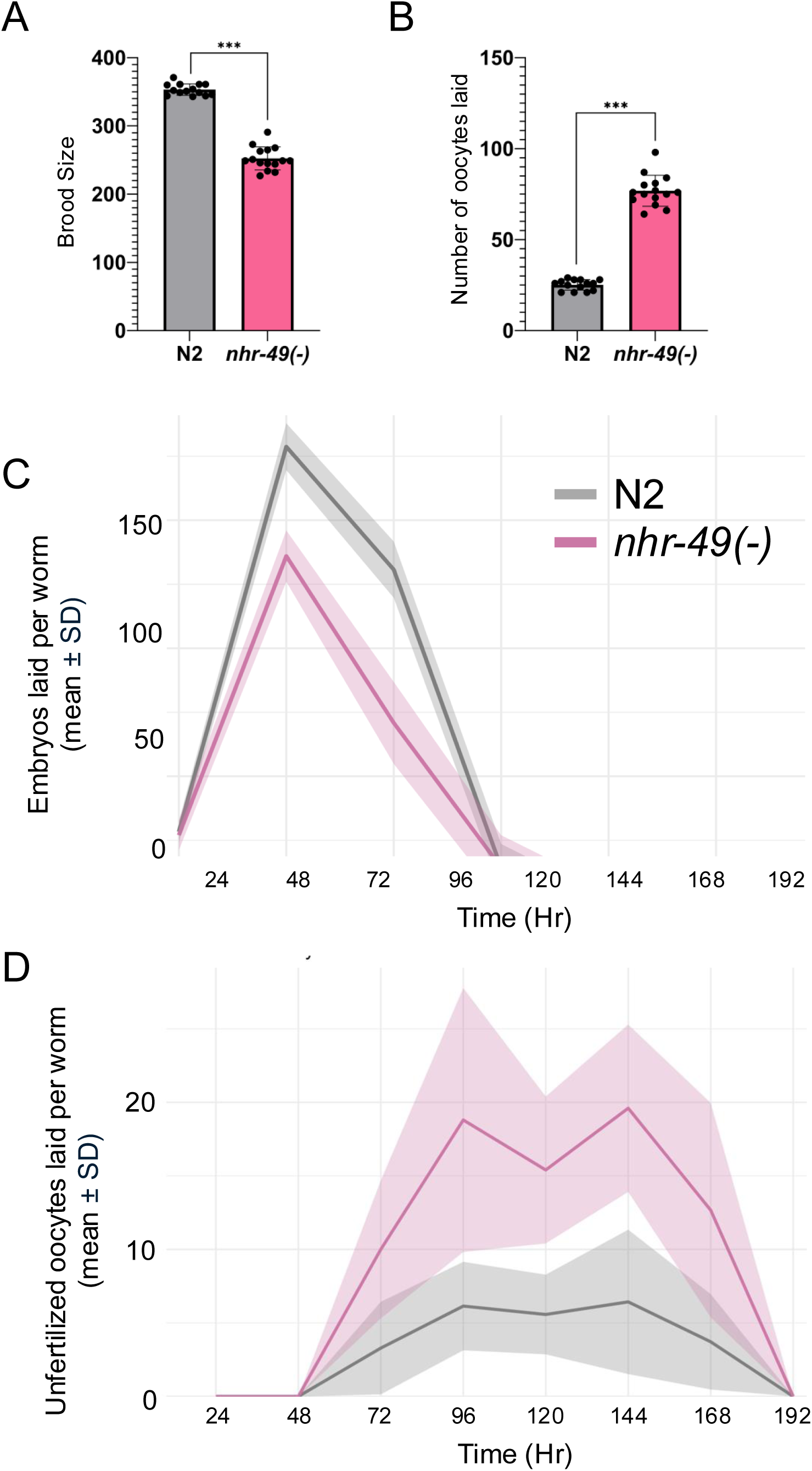
NHR-49 is required for the retention of unfertilized oocytes in reproductive and post-reproductive hermaphrodites. **(A)** Bar graphs showing mean brood size of N2 (wild-type) and *nhr-49(gk405)* mutants. **(B)** Bar graphs showing the mean number of unfertilized oocytes laid by N2 (wild-type) and *nhr-49(gk405)* mutants on day 6 of adulthood. **(C-D)** Line graphs showing the number of embryos (C) and unfertilized oocytes (mean±SD) (D) laid by *N2* and *nhr-49(gk405)* worms throughout their reproductive span. Mean values were calculated based on the number of embryos or oocytes laid by at least 14 worms from two independent replicates. Embryo and unfertilized oocyte counts were recorded starting from the L4 larval stage.

**Figure S2.**
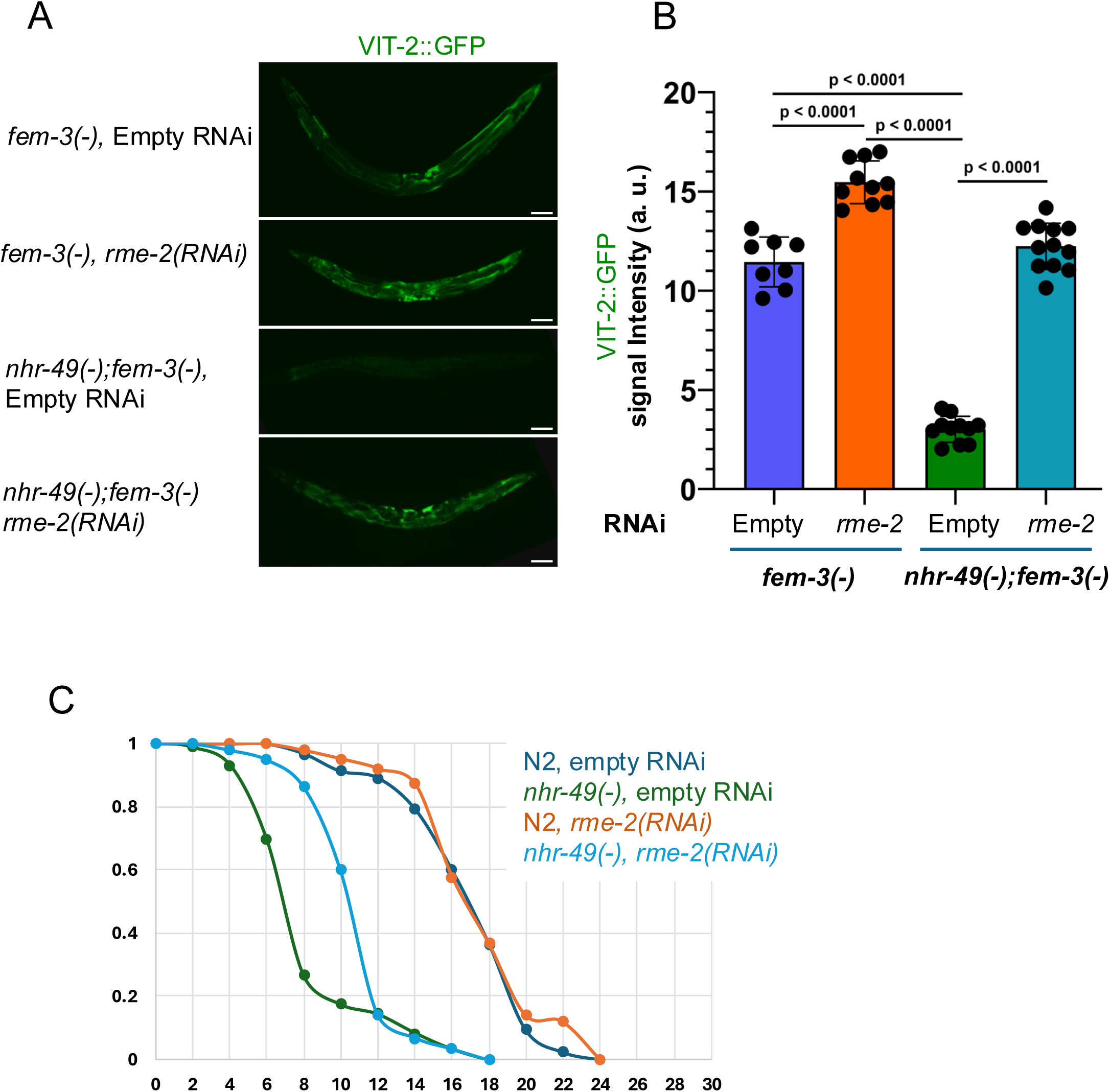
Lifespan shortening and fat loss in *nhr-49* mutants are partially dependent on RME-2-mediated yolk transport into oocytes. **(A)** Representative images of VIT-2::GFP (YP170::GFP) ^42^expression in *fem-3(e1996);nhr-49(gk405)* worms with and without *rme-2* RNAi. **(B)** Bar graph showing fluorescence intensity in the indicated strains on day 1 of adulthood. Fluorescence intensity is shown as the mean intensity (N >10) for the indicated genotype with or without *rme-2* RNAi. Two-tailed P values were computed using an unpaired t-test. **(E)** Survival curves for N2 (Wildtype), and *nhr-49(gk405)* mutants treated with control RNAi (EV) and *rme-2* RNAi from one representative experiment. Quantitative data and Kaplan–Meier statistics are provided in table 2.

**Figure S3.**
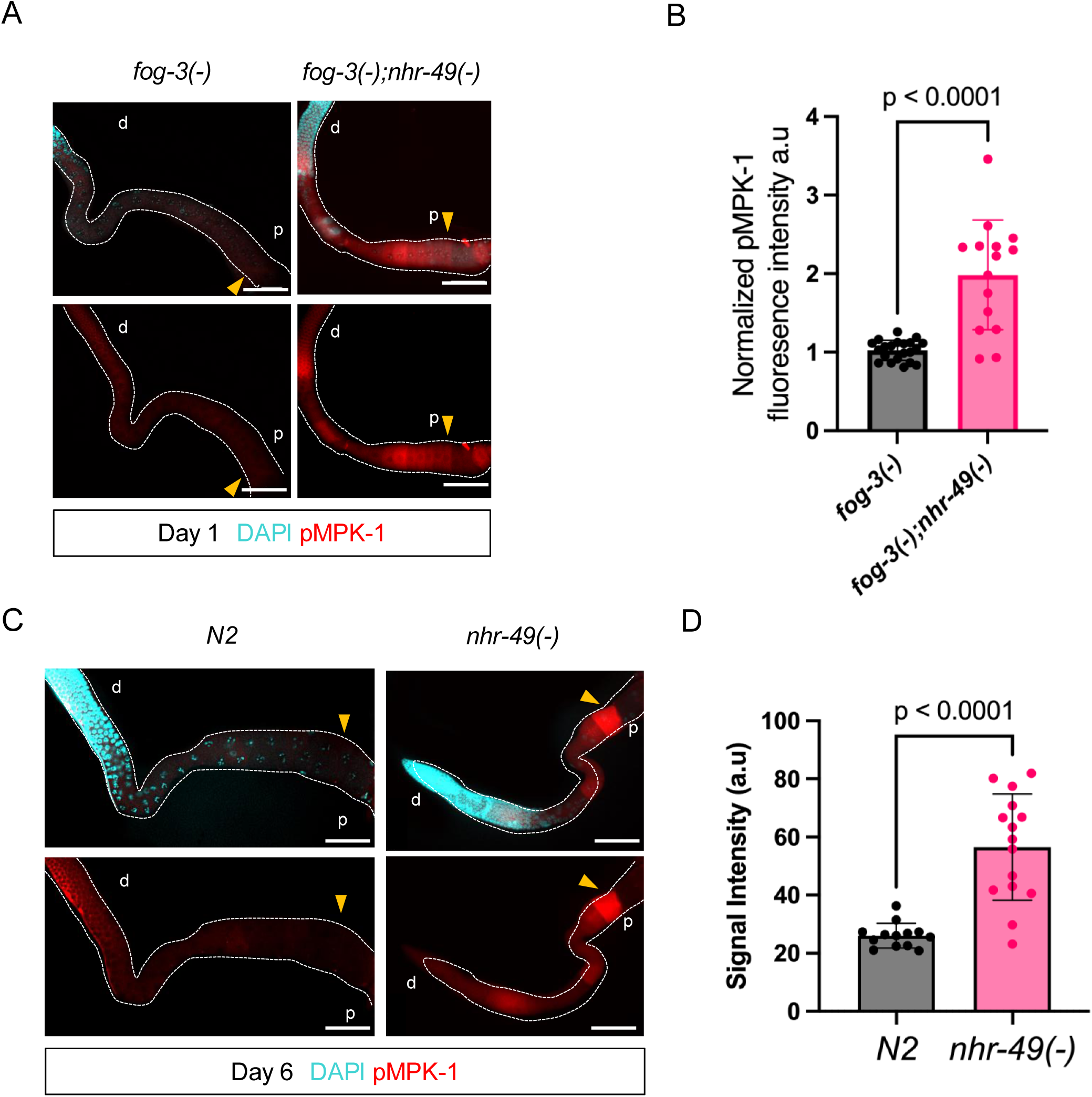
NHR-49 negatively regulates oocyte activation in the proximal gonad of feminized mutants. **(A)** Representative images of dpMPK-1 staining (red) of dissected gonads from day-1 adult feminized *fog-3(q443)* and *fog-3(q443);nhr-49(gk405)* double mutants. The position of the oocyte most proximal to the spermatheca in each image is marked by a yellow arrowhead. **(B)** Bar graph showing dpMPK-1 mean signal intensity in the proximal oocytes of the strains indicated in (A). Two-tailed p-values were computed using an unpaired t-test. **(C)** Representative images of dpMPK-1 staining of dissected gonads from post-reproductive N2 (wild-type) and *nhr-49(gk405)* mutants at day 6 adulthood. The position of the oocyte most proximal to the spermatheca in each image is marked by a yellow arrowhead. **(D)** Bar graph showing dpMPK-1 mean signal intensity in the proximal oocytes of the strains indicated in (C). Two-tailed p-value were computed using an unpaired t-test.

**Figure S4.**
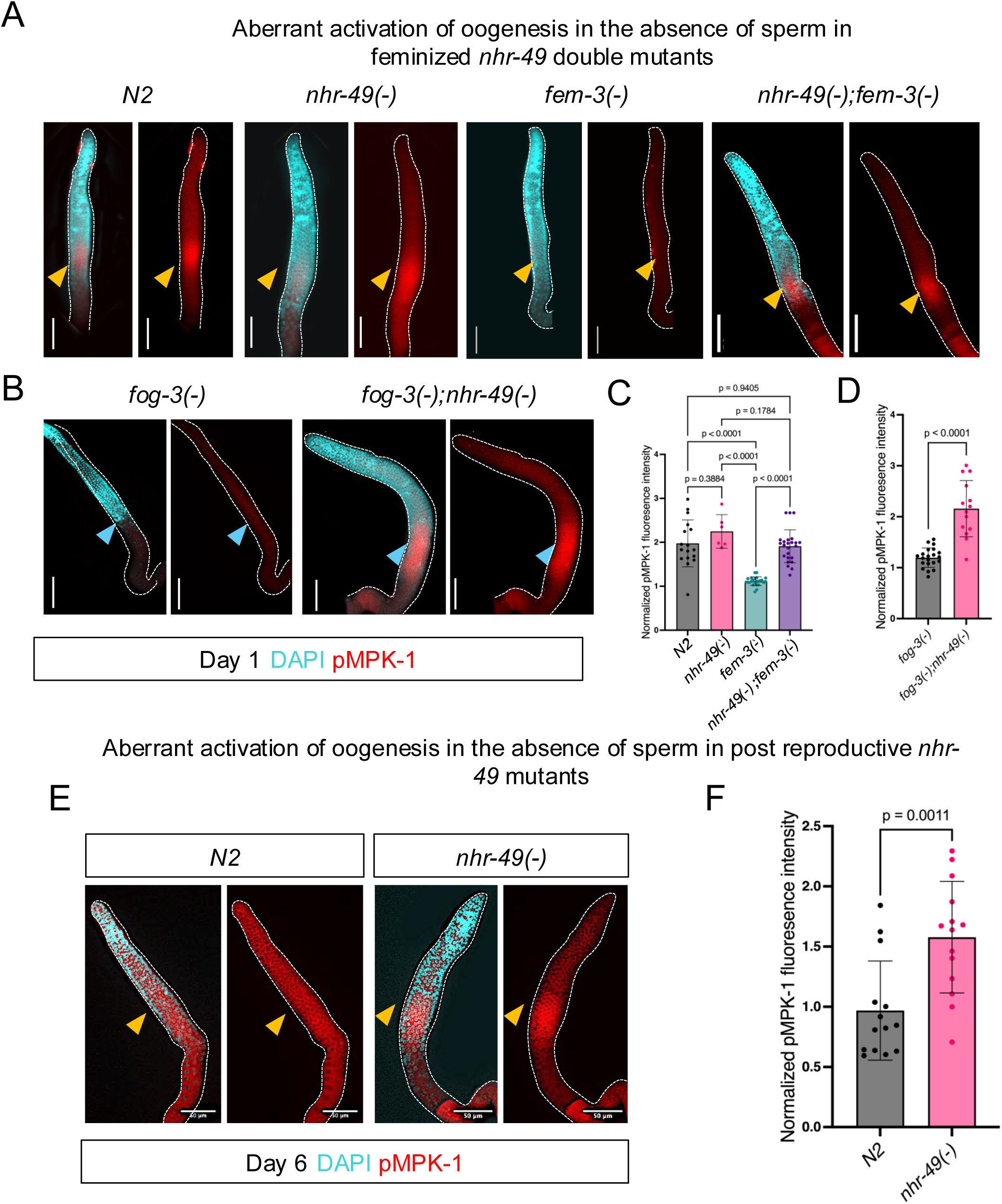
NHR-49 promotes germline MPK-1 activation in feminized and post-reproductive animals. **(A-B)** Representative images of dpMPK-1 staining (red) in dissected gonads from day 1 adult N2 (wild-type), *nhr-49(gk405)*, *fem-3(e1996)*, *fem-3(e1996);nhr-49(gk405)*, *fog-3(q443)*, and *fog-3(q443);nhr-49(gk405)*. DAPI (cyan) marks germ cell nuclei. The position of the oocyte most proximal to the spermatheca in each image is marked by a yellow or arrowhead. **(C-D)** Bar graph showing dpMPK-1 signal intensity in the proximal oocytes of the indicated strains. Statistical comparisons were performed using one-way analysis of variance (ANOVA) followed by Tukey’s multiple comparisons test. More than 45 worms from 2 independent replicates per genotype were analyzed. (E) Representative images of dpMPK-1 staining in dissected gonads from day 6 N2 and *nhr-49(gk405)* worms. Scale bars = 50 µm. (F) Bar graph showing dpMPK-1 signal intensity in the proximal oocytes of **day 6 post-reproductive adults** of the indicated strains. Statistical comparisons were performed using one-way analysis of variance (ANOVA) followed by Tukey’s multiple comparisons test. More than 45 worms from 2 independent replicates per genotype were analyzed. **Panel (D) and (F) were analyzed using unpaired two-tailed t-tests. Bars represent mean ± SD.**

**Figure S5.**
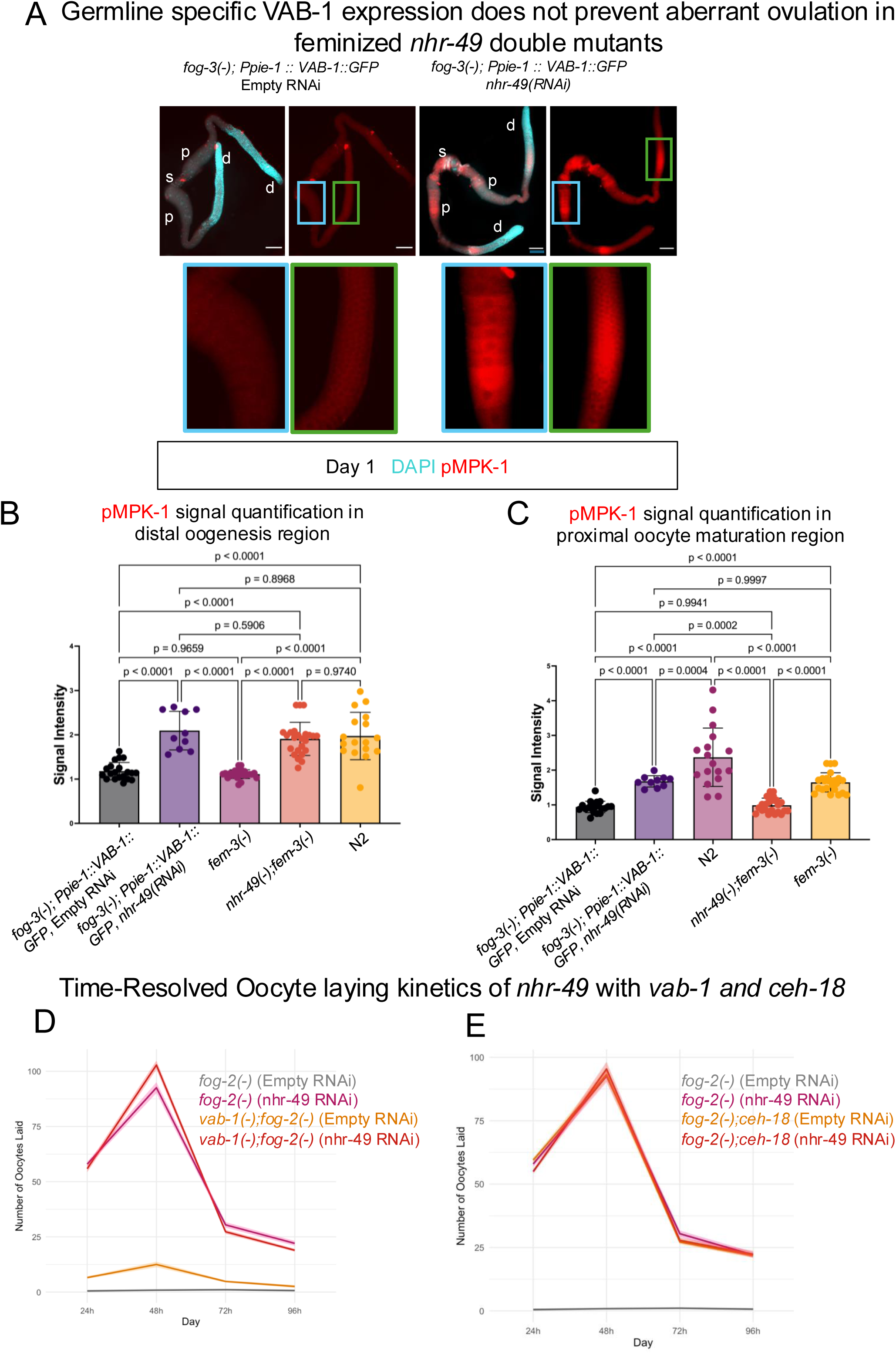
NHR-49 likely regulates oocyte activation primarily through somatic tissues. **(A-B)** Bar graphs showing mean dpMPK-1 signal intensities from immunostained gonads of *fog-3(q443);Ppie-1::VAB-1::GFP* worms, in which *vab-1* is driven by a germline-specific promoter *pie-1*. Quantification of signal intensities in the distal oogenesis region (A) and in the proximal oocyte maturation region (B). These worms still showed aberrant dpMPK-1 expression upon *nhr-49* RNAi in both distal and proximal regions. Comparisons were statistically tested by one-way analysis of variance (ANOVA) followed by multiple comparisons with Tukey’s correction, based on more than 45 worms from two independent replicates per genotype. **(C)** Representative images of dpMPK-1 staining (red) in dissected gonads of *fog-3(q443);Ppie-1::VAB-1::GFP* animals. Animals were treated with control (EV) or *nhr-49* RNAi. DAPI staining (blue) was used to stain germ cell nuclei. Blue box indicates proximal maturation region, and the green box indicates distal oogenesis region. (D) Line graphs showing the number of unfertilized oocytes (mean±SD) laid by *fog-2(q71), vab-1(dx31);fog-2(q71)* (E) *fog-2(q71), fog-2(q71);ceh-18(mg57)* animals treated with control (EV) or *nhr-49* RNAi. Mean values were calculated based on the number of embryos or oocytes laid by at least 15 worms from two independent replicates. Embryo and unfertilized oocyte counts were recorded starting from the L4 larval stage.

**Figure S6.**
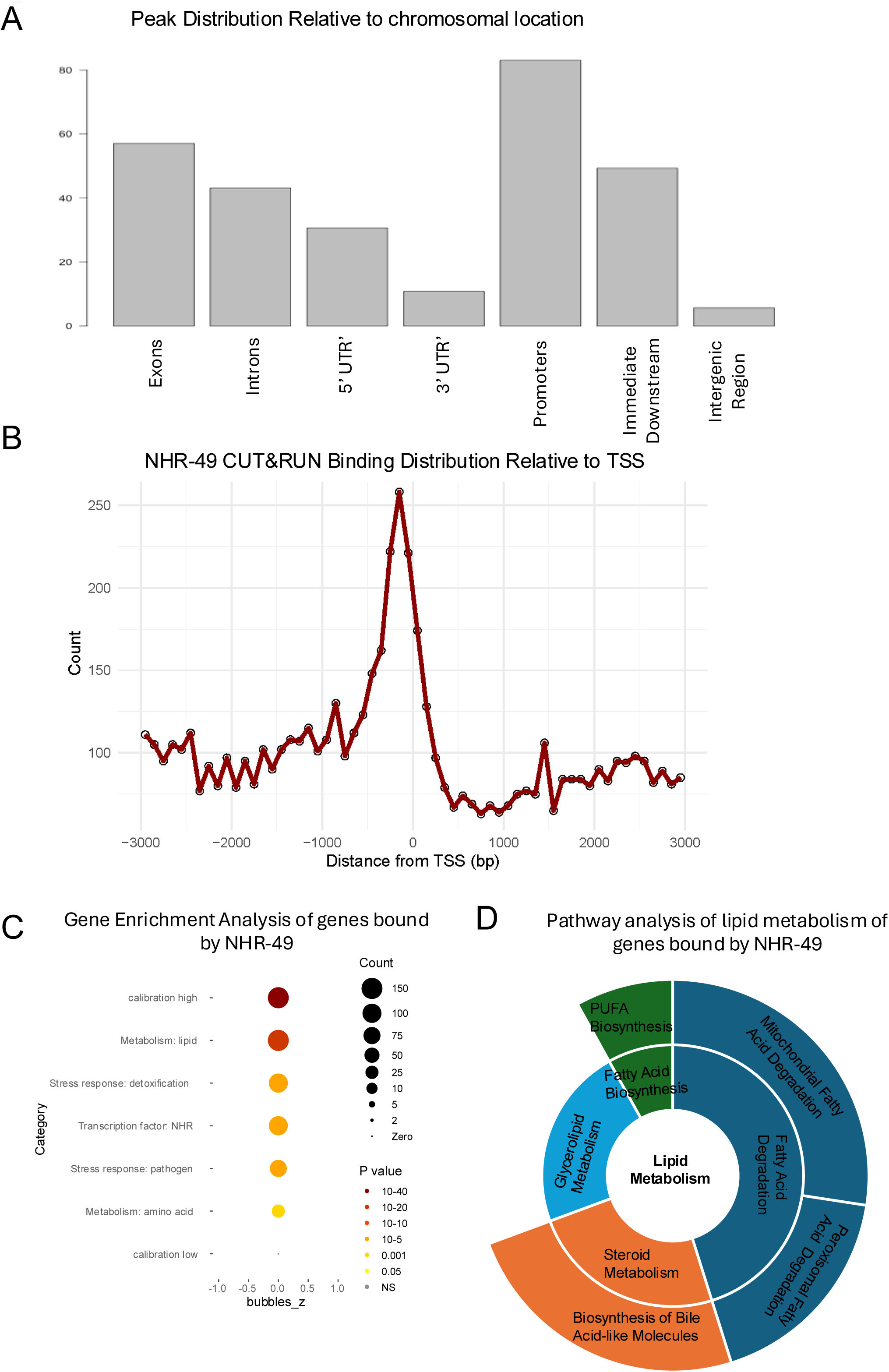
Genome-wide binding profile of NHR-49::HA revealed by CUT&RUN. **(A)** Distribution of NHR-49 binding peaks across genomic features. The majority of NHR-49 binding is localized to promoter regions. **(B)** NHR-49 binding peak distribution relative to transcription start sites (TSS). The majority of NHR-49 binding peaks are located within 1kb upstream of the TSS. **(C)** WormCat gene set enrichment analysis of NHR-49 binding targets. Dot size represents the number of genes; color indicates adjusted *p*-value. **(D)** KEGG pathway analysis of lipid metabolism genes associated with NHR-49 CUT&RUN binding peaks. Sunburst plot shows enriched lipid metabolic pathways, with the inner ring representing broader categories (e.g., fatty acid metabolism, steroid metabolism), and the outer ring representing more specific sub-pathways.

**Fig S7.**
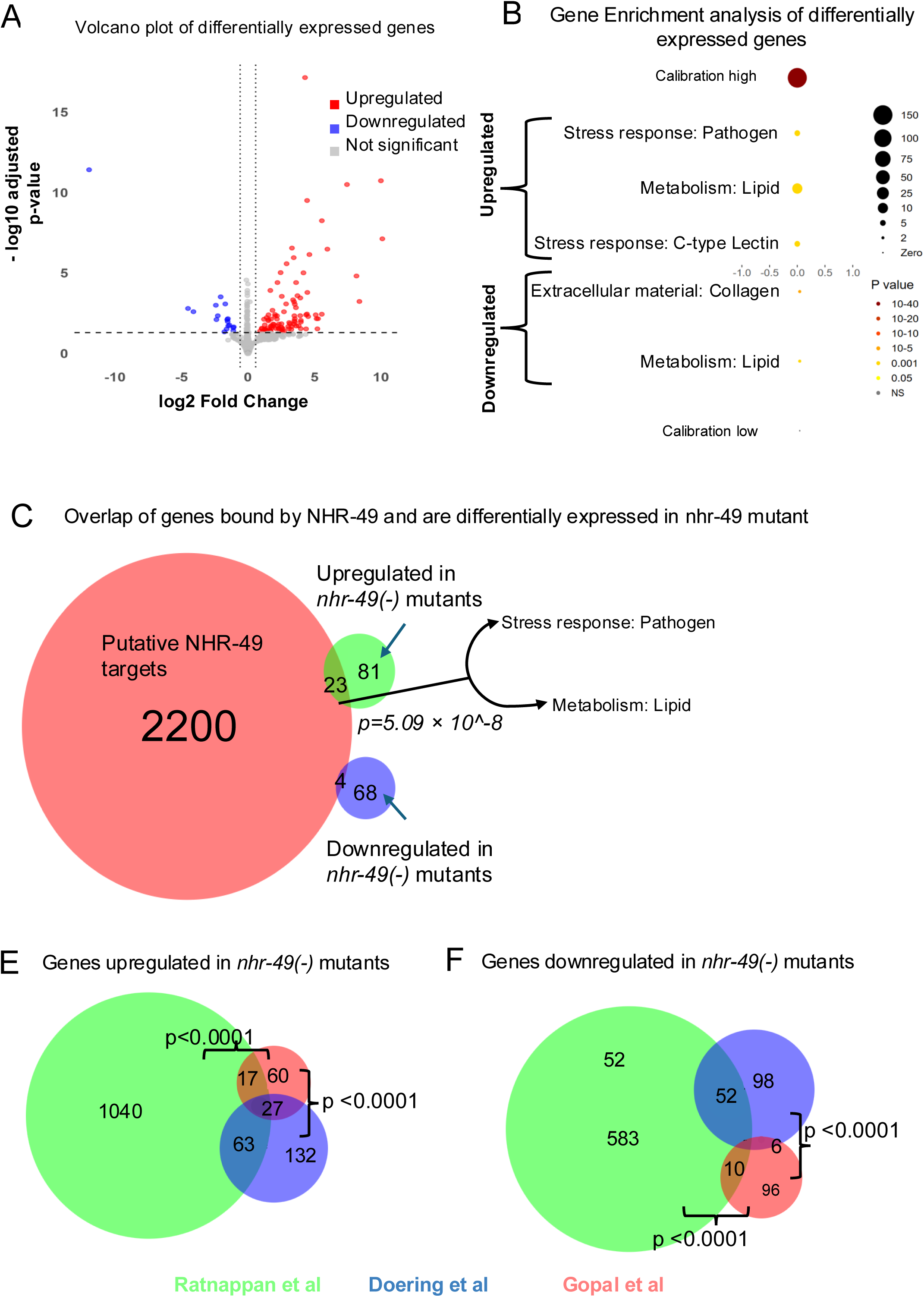
NHR-49 binding targets overlap with genes differentially expressed in *nhr-49* mutants and across published whole-worm transcriptomes. **(A)** Volcano plot showing differential gene expression in *nhr-49(gk405)* mutants compared to N2 (wild-type) animals, based on whole-worm RNA-seq analyzed using DESeq2. Significantly upregulated (red) and downregulated (blue) genes are indicated (adjusted *p* < 0.05, log₂FC > 0.58). **(B)** Gene set enrichment analysis of genes upregulated (top) and downregulated (bottom) in *nhr-49(gk405)* mutants. Dot size represents the number of genes; color indicates adjusted *p*-value. **(C)** Venn diagram showing overlap between NHR-49 CUT&RUN binding targets (red) and genes upregulated (green) or downregulated (blue) in *nhr-49(gk405)* mutants. *p*-value calculated using Fisher’s exact test. **(D)** Gene set enrichment analysis of the genes overlapping between NHR-49 CUT&RUN binding targets and differentially expressed gene in *nhr-49(gk405)* mutants. **(E, F)** Venn diagrams comparing genes upregulated (E) or downregulated (F) in this study with previously published whole-worm transcriptomes from Ratnappan et al^33^ (2014) and Doering et al^19^ (2021).

## References

1. Stearns, S.C. (1989). Trade-Offs in Life-History Evolution. Funct. Ecol. 3, 259–268. 10.2307/2389364.

2. Della Torre, S., Benedusi, V., Fontana, R., and Maggi, A. (2014). Energy metabolism and fertility: a balance preserved for female health. Nat. Rev. Endocrinol. 10, 13–23. 10.1038/nrendo.2013.203.

3. Mircea, C.N., Lujan, M.E., and Pierson, R.A. (2007). Metabolic fuel and clinical implications for female reproduction. J. Obstet. Gynaecol. Can. JOGC J. Obstet. Gynecol. Can. JOGC 29, 887–902. 10.1016/S1701-2163(16)32661-5.

4. Lopez, A.L., Chen, J., Joo, H.-J., Drake, M., Shidate, M., Kseib, C., and Arur, S. (2013). DAF-2 and ERK Couple Nutrient Availability to Meiotic Progression during *Caenorhabditis elegans* Oogenesis. Dev. Cell 27, 227–240. 10.1016/j.devcel.2013.09.008.

5. Sahu, S., and Armstrong, A.R. (2024). Translation components in adult Drosophila melanogaster adipocytes regulate the ovarian germline stem cell lineage. Preprint at bioRxiv, 10.1101/2024.08.31.610632 https://doi.org/10.1101/2024.08.31.610632.

6. Thondamal, M., Witting, M., Schmitt-Kopplin, P., and Aguilaniu, H. (2014). Steroid hormone signalling links reproduction to lifespan in dietary-restricted Caenorhabditis elegans. Nat. Commun. 5, 4879. 10.1038/ncomms5879.

7. Templeman, N.M., and Murphy, C.T. (2018). Regulation of reproduction and longevity by nutrient-sensing pathways. J. Cell Biol. 217, 93–106. 10.1083/jcb.201707168.

8. Cao, W., and Pocock, R. (2022). Mechanisms of germ cell survival and plasticity in Caenorhabditis elegans. Biochem. Soc. Trans. 50, 1517–1526. 10.1042/BST20220878.

9. Brenner, S. (1974). The genetics of Caenorhabditis elegans. Genetics 77, 71–94. 10.1093/genetics/77.1.71.

10. Pazdernik, N., and Schedl, T. (2013). Introduction to Germ Cell Development in C. elegans. Adv. Exp. Med. Biol. 757, 1–16. 10.1007/978-1-4614-4015-4_1.

11. Kimble, J., and Crittenden, S.L. (2005). Germline proliferation and its control. WormBook Online Rev. C Elegans Biol., 1–14. 10.1895/wormbook.1.13.1.

12. Davis, G.M., Hipwell, H., and Boag, P.R. (2023). Oogenesis in Caenorhabditis elegans. Sex. Dev. 17, 73–83. 10.1159/000531019.

13. Hodgkin, J. (1986). Sex Determination in the Nematode C. ELEGANS: Analysis of tra-3 Suppressors and Characterization of fem Genes. Genetics 114, 15–52. 10.1093/genetics/114.1.15.

14. Schedl, T., and Kimble, J. (1988). Fog-2, a Germ-Line-Specific Sex Determination Gene Required for Hermaphrodite Spermatogenesis in Caenorhabditis Elegans. Genetics 119, 43–61. 10.1093/genetics/119.1.43.

15. Cheng, H., Govindan, J.A., and Greenstein, D. (2008). Regulated trafficking of the MSP/Eph receptor during oocyte meiotic maturation in C. elegans. Curr. Biol. CB 18, 705–714. 10.1016/j.cub.2008.04.043.

16. Miller, M.A., Nguyen, V.Q., Lee, M.-H., Kosinski, M., Schedl, T., Caprioli, R.M., and Greenstein, D. (2001). A Sperm Cytoskeletal Protein That Signals Oocyte Meiotic Maturation and Ovulation. Science 291, 2144–2147. 10.1126/science.1057586.

17. Miller, M.A., Ruest, P.J., Kosinski, M., Hanks, S.K., and Greenstein, D. (2003). An Eph receptor sperm-sensing control mechanism for oocyte meiotic maturation in Caenorhabditis elegans. Genes Dev. 17, 187–200. 10.1101/gad.1028303.

18. Chaturbedi, A., and Lee, S.S. (2023). Different gametogenesis states uniquely impact longevity in Caenorhabditis elegans. Preprint at bioRxiv, 10.1101/2023.06.13.544885 https://doi.org/10.1101/2023.06.13.544885.

19. Doering, K.R.S., Ermakova, G., and Taubert, S. (2023). Nuclear hormone receptor NHR-49 is an essential regulator of stress resilience and healthy aging in Caenorhabditis elegans. Front. Physiol. 14, 1241591. 10.3389/fphys.2023.1241591.

20. Scholtes, C., and Giguère, V. (2022). Transcriptional control of energy metabolism by nuclear receptors. Nat. Rev. Mol. Cell Biol. 23, 750–770. 10.1038/s41580-022-00486-7.

21. Sever, R., and Glass, C.K. (2013). Signaling by Nuclear Receptors. Cold Spring Harb. Perspect. Biol. 5, a016709. 10.1101/cshperspect.a016709.

22. Taubert, S., Van Gilst, M.R., Hansen, M., and Yamamoto, K.R. (2006). A Mediator subunit, MDT-15, integrates regulation of fatty acid metabolism by NHR-49-dependent and - independent pathways in C. elegans. Genes Dev. 20, 1137–1149. 10.1101/gad.1395406.

23. Van Gilst, M.R., Hadjivassiliou, H., Jolly, A., and Yamamoto, K.R. (2005). Nuclear hormone receptor NHR-49 controls fat consumption and fatty acid composition in C. elegans. PLoS Biol. 3, e53. 10.1371/journal.pbio.0030053.

24. Goh, G.Y.S., Winter, J.J., Bhanshali, F., Doering, K.R.S., Lai, R., Lee, K., Veal, E.A., and Taubert, S. (2018). NHR-49/HNF4 integrates regulation of fatty acid metabolism with a protective transcriptional response to oxidative stress and fasting. Aging Cell 17, e12743. 10.1111/acel.12743.

25. Doering, K.R., Cheng, X., Milburn, L., Ratnappan, R., Ghazi, A., Miller, D.L., and Taubert, S. (2022). Nuclear hormone receptor NHR-49 acts in parallel with HIF-1 to promote hypoxia adaptation in Caenorhabditis elegans. eLife 11, e67911. 10.7554/eLife.67911.

26. Zeng, L., Li, X., Preusch, C.B., He, G.J., Xu, N., Cheung, T.H., Qu, J., and Mak, H.Y. (2021). Nuclear receptors NHR-49 and NHR-79 promote peroxisome proliferation to compensate for aldehyde dehydrogenase deficiency in C. elegans. PLOS Genet. 17, e1009635. 10.1371/journal.pgen.1009635.

27. Kwon, S., Park, K.-S., and Yoon, K. (2024). Regulator of Lipid Metabolism NHR-49 Mediates Pathogen Avoidance through Precise Control of Neuronal Activity. Cells 13, 978. 10.3390/cells13110978.

28. Dasgupta, M., Shashikanth, M., Sandhu, A., De, A., Javed, S., and Singh, V. (2020). NHR-49 Transcription Factor Regulates Immunometabolic Response and Survival of Caenorhabditis elegans during Enterococcus faecalis Infection. Infect. Immun. 88. 10.1128/IAI.00130-20.

29. Naim, N., Amrit, F.R.G., Ratnappan, R., DelBuono, N., Loose, J.A., and Ghazi, A. (2021). Cell nonautonomous roles of NHR-49 in promoting longevity and innate immunity. Aging Cell 20, e13413. 10.1111/acel.13413.

30. Eustice, M., Konzman, D., Reece, J.M., Ghosh, S., Alston, J., Hansen, T., Golden, A., Bond, M.R., Abramowitz, L.K., and Hanover, J.A. (2022). Nutrient sensing pathways regulating adult reproductive diapause in C. elegans. PloS One 17, e0274076. 10.1371/journal.pone.0274076.

31. Seidel, H.S., and Kimble, J. (2011). The oogenic germline starvation response in C. elegans. PloS One 6, e28074. 10.1371/journal.pone.0028074.

32. Heestand, B., Simon, M., Frenk, S., Titov, D., and Ahmed, S. (2018). Transgenerational Sterility of Piwi Mutants Represents a Dynamic Form of Adult Reproductive Diapause. Cell Rep. 23, 156–171. 10.1016/j.celrep.2018.03.015.

33. Ratnappan, R., Amrit, F.R.G., Chen, S.-W., Gill, H., Holden, K., Ward, J., Yamamoto, K.R., Olsen, C.P., and Ghazi, A. (2014). Germline signals deploy NHR-49 to modulate fatty-acid β-oxidation and desaturation in somatic tissues of C. elegans. PLoS Genet. 10, e1004829. 10.1371/journal.pgen.1004829.

34. Nuclear hormone receptor NHR-49 acts in parallel with HIF-1 to promote hypoxia adaptation in Caenorhabditis elegans | eLife https://elifesciences.org/articles/67911.

35. Schedl, T., and Kimble, J. (1988). fog-2, a germ-line-specific sex determination gene required for hermaphrodite spermatogenesis in Caenorhabditis elegans. Genetics 119, 43–61. 10.1093/genetics/119.1.43.

36. Scharf, A., Pohl, F., Egan, B.M., Kocsisova, Z., and Kornfeld, K. (2021). Reproductive Aging in Caenorhabditis elegans: From Molecules to Ecology. Front. Cell Dev. Biol. 9, 718522. 10.3389/fcell.2021.718522.

37. Rapid Lipid Quantification in Caenorhabditis elegans by Oil Red O and Nile Red Staining https://en.bio-protocol.org/en/bpdetail?id=4340&type=0.

38. Hou, N.S., and Taubert, S. (2012). Function and Regulation of Lipid Biology in Caenorhabditis elegans Aging. Front. Physiol. 3. 10.3389/fphys.2012.00143.

39. Mullaney, B.C., and Ashrafi, K. (2009). C. elegans Fat Storage and Metabolic Regulation. Biochim. Biophys. Acta 1791, 474–478. 10.1016/j.bbalip.2008.12.013.

40. Evolutionarily related host and microbial pathways regulate fat desaturation in C. elegans | Nature Communications https://www.nature.com/articles/s41467-024-45782-2.

41. Balklava, Z., Pant, S., Fares, H., and Grant, B.D. (2007). Genome-wide analysis identifies a general requirement for polarity proteins in endocytic traffic. Nat. Cell Biol. 9, 1066–1073. 10.1038/ncb1627.

42. Grant, B., and Hirsh, D. (1999). Receptor-mediated Endocytosis in the Caenorhabditis elegans Oocyte. Mol. Biol. Cell 10, 4311–4326. 10.1091/mbc.10.12.4311.

43. Bohnert, K.A., and Kenyon, C. (2017). A lysosomal switch triggers proteostasis renewal in the immortal C. elegans germ lineage. Nature 551, 629–633. 10.1038/nature24620.

44. Kocsisova, Z., Mohammad, A., Kornfeld, K., and Schedl, T. (2018). Cell Cycle Analysis in the C. elegans Germline with the Thymidine Analog EdU. J. Vis. Exp. JoVE, 58339. 10.3791/58339.

45. Watterson, A., Tatge, L., Wajahat, N., Arneaud, S.L.B., Solano Fonseca, R., Beheshti, S.T., Metang, P., Mihelakis, M., Zuurbier, K.R., Corley, C.D., et al. (2022). Intracellular lipid surveillance by small G protein geranylgeranylation. Nature 605, 736–740. 10.1038/s41586-022-04729-7.

46. Angelo, G., and Van Gilst, M.R. (2009). Starvation Protects Germline Stem Cells and Extends Reproductive Longevity in *C. elegans*. Science 326, 954–958. 10.1126/science.1178343.

47. Yang, Y., Han, S.M., and Miller, M.A. (2010). MSP Hormonal Control of the Oocyte MAP Kinase Cascade and Reactive Oxygen Species Signaling. Dev. Biol. 342, 96–107. 10.1016/j.ydbio.2010.03.026.

48. Construction of a germline-specific RNAi tool in C. elegans https://www.ncbi.nlm.nih.gov/pmc/articles/PMC6382888/.

49. Holdorf, A.D., Higgins, D.P., Hart, A.C., Boag, P.R., Pazour, G.J., Walhout, A.J.M., and Walker, A.K. (2020). WormCat: An Online Tool for Annotation and Visualization of Caenorhabditis elegans Genome-Scale Data. Genetics 214, 279–294. 10.1534/genetics.119.302919.

50. Govindan, J.A., Cheng, H., Harris, J.E., and Greenstein, D. (2006). Gαo/i and Gαs Signaling Function in Parallel with the MSP/Eph Receptor to Control Meiotic Diapause in C. elegans. Curr. Biol. 16, 1257–1268. 10.1016/j.cub.2006.05.020.

51. Martin, M. (2011). Cutadapt removes adapter sequences from high-throughput sequencing reads. EMBnet.journal 17, 10–12. 10.14806/ej.17.1.200.

52. Dobin, A., Davis, C.A., Schlesinger, F., Drenkow, J., Zaleski, C., Jha, S., Batut, P., Chaisson, M., and Gingeras, T.R. (2013). STAR: ultrafast universal RNA-seq aligner. BIOINFORMATICS 29, 15–21. 10.1093/bioinformatics/bts635.

53. Love, M.I., Huber, W., and Anders, S. (2014). Moderated estimation of fold change and dispersion for RNA-seq data with DESeq2. Genome Biol. 15, 550. 10.1186/s13059-014-0550-8.

54. Hulsen, T., de Vlieg, J., and Alkema, W. (2008). BioVenn – a web application for the comparison and visualization of biological lists using area-proportional Venn diagrams. BMC Genomics 9, 488. 10.1186/1471-2164-9-488.

55. Emerson, F.J., and Lee, S.S. (2022). CUT&RUN for Chromatin Profiling in Caenorhabditis elegans. Curr. Protoc. 2, e445. 10.1002/cpz1.445.

56. Skene, P.J., Henikoff, J.G., and Henikoff, S. (2018). Targeted in situ genome-wide profiling with high efficiency for low cell numbers. Nat. Protoc. 13, 1006–1019. 10.1038/nprot.2018.015.

57. Skene, P.J., and Henikoff, S. (2017). An efficient targeted nuclease strategy for high-resolution mapping of DNA binding sites. eLife 6, e21856. 10.7554/eLife.21856.

58. Li, C.-L., Pu, M., Wang, W., Chaturbedi, A., Emerson, F.J., and Lee, S.S. (2021). Region-specific H3K9me3 gain in aged somatic tissues in Caenorhabditis elegans. PLOS Genet. 17, e1009432. 10.1371/journal.pgen.1009432.

59. Emerson, F.J., Chiu, C., Lin, L.Y., Riedel, C.G., Zhu, M., and Lee, S.S. (2024). The chromatin factors SET-26 and HCF-1 oppose the histone deacetylase HDA-1 in longevity and gene regulation in C. elegans. Nat. Commun. 15, 2320. 10.1038/s41467-024-46510-6.

60. Zhang, Y., Liu, T., Meyer, C.A., Eeckhoute, J., Johnson, D.S., Bernstein, B.E., Nusbaum, C., Myers, R.M., Brown, M., Li, W., et al. (2008). Model-based Analysis of ChIP-Seq (MACS). Genome Biol. 9, R137. 10.1186/gb-2008-9-9-r137.

